# Extended correlation functions for spatial analysis of multiplex imaging data

**DOI:** 10.1101/2023.06.20.545678

**Authors:** Joshua A. Bull, Eoghan J. Mulholland, Simon J. Leedham, Helen M. Byrne

## Abstract

Imaging platforms for generating highly multiplexed histological images are being continually developed and improved. Significant improvements have also been made in the accuracy of methods for automated cell segmentation and classification. However, less attention has focussed on the quantification and analysis of the resulting point clouds which describe the spatial coordinates of individual cells. We focus here on a particular spatial statistical method, the cross-pair correlation function (cross-PCF), which can identify positive and negative spatial correlation between cells across a range of length scales. However, limitations of the cross-PCF hinder its widespread application to multiplexed histology. For example, it can only consider relations between pairs of cells, and cells must be classified using discrete categorical labels (rather than labelling continuous labels such as stain intensity).

In this paper, we present three extensions to the cross-PCF which address these limitations and permit more detailed analysis of multiplex images: Topographical Correlation Maps (TCMs) can visualise local clustering and exclusion between cells; Neighbourhood Correlation Functions (NCFs) can identify colocalisation of two or more cell types; and weighted-PCFs (wPCFs) describe spatial correlation between points with continuous (rather than discrete) labels. We apply the extended PCFs to synthetic and biological datasets in order to demonstrate the insight that they can generate.

**Impact statement:** This paper introduces three methods for performing spatial analysis on multiplex digital pathology images. We apply the methods to synthetic datasets and regions of interest from a murine colorectal carcinoma, in order to illustrate their relative strengths and weaknesses. We note that these methods have wider application to marked point pattern data from other sources.

## Introduction

The move to digital pathology is revolutionising the way in which histological samples are processed, viewed and analysed. Until recently, pathology was restricted to expert manual assessment of hematoxylin and eosin (H&E) and immunohistochemistry (IHC) slides stained with a small number of coloured dyes. Multiplex modalities now enable digital visualisation of whole slide images (WSIs), stained with relatively large numbers of markers, at sub-micrometer resolution. Digital pathology slides can be generated using a variety of methods, including multiplex immunohistochemistry, imaging mass cytometry (IMC), co-detection by indexing (CODEX/Phenocycler), and multiplexed ion beam imaging (MIBI)^(1–4)^. These platforms can generate images with 50 or more cellular markers (see, e.g., ^(5)^). As the number of cell types discernible in a multiplex image increases, simply viewing an image can be challenging because of the difficulty in choosing a unique colouring for each cell marker. Additionally, existing statistical methods struggle to exploit the full range of spatial information contained within the data, with analysis dominated by nonspatial metrics such as cell counts or basic spatial metrics such as mean intercellular distances which do not account for the wider spatial context within an image. While the methodology underlying different imaging technologies may vary, the images they generate all encode high resolution information about the spatial location of multiple cell markers. As such, computational methods developed to analyse cell locations generated from one multiplex modality can be applied straightforwardly to data generated from another.

State-of-the-art pipelines for the statistical analysis of multiplex images typically involve at least two preprocessing steps: *cell segmentation*, in which the boundaries of individual cells are identified, and *cell classification*, in which cells are assigned to categories based on the panel of markers used for image generation^(6–9)^. The accuracy of cell segmentation has improved significantly in recent years, driven primarily by advances in artificial intelligence (AI) based approaches for cell detection^(10,11)^. Many of these methods can be accessed via open source digital pathology platforms such as Qupath^(12)^ or MCMICRO^(13)^, commercial tools such as HALO (indicalab.com/halo) and Visiopharm (visiopharm.com), and standalone software such as Deepcell^(10)^ and Cellpose^(11)^. By contrast, there are fewer tools for cell classification, due perhaps to variation in the panels used for a given study. Existing tools are typically iterative and semi-supervised^(6,7,14)^.

The above improvements in preprocessing digital pathology slides are increasing the demand for methods that can describe and quantify the spatial information contained within multiplex images. Such information is important because there is increasing evidence that physical contact can alter cell behaviours and drive disease progression. For example, the formation of tumour microenvironment of metastasis (or ‘TMEM’) doorways is implicated in the metastasis of cancer stem cells^(28,29)^. TMEMs form when a Mena^Hi^ tumour cell, a macrophage, and an endothelial cell come into physical contact on the surface of a blood vessel^(30)^. This three-way spatial interaction enables tumour cells first to intravasate and then to metastasise to other parts of the body, and has also been implicated in cancer cell acquisition of a stem-like phenotype^(30)^. Other biological effects which manifest in altered spatial interactions include clustering of immune cells and alveolar progenitor cells in the lungs during COVID-19 progression^(7)^, and the formation of distinct cellular neighbourhoods which drive antitumoral immune responses in the invasive front of colorectal cancer. For example, neighbourhoods which are rich in both granuloctyes and PD-1+CD4+ T cells correlate positively with patient survival^(31)^. While spatially averaged statistics, such as cell counts, can be readily calculated from segmented and classified images, describing and quantifying the spatial organisation of cell types requires more complex analytical tools.

One promising approach for exploiting the spatial structure of multiplex images is AI and machine learning, which learns to identify those regions of an image which are most strongly associated with clinical features such as patient prognosis and disease status^(32,33)^. Machine learning approaches include convolutional neural networks (CNNs), generative adversarial networks (GANs) and transformers. They have been used to perform a range of tasks, such as automatic identification of informative regions in whole slide images (WSIs)^(34)^, segmentation of Ductal Carcinoma In Situ (DCIS)^(35)^, and prediction of molecular signatures from tissue morphology^(36)^. However, such machine learning methods typically require large training datasets and it can be difficult to understand or interpret their predictions. Further, machine learning methods usually require the same marker combinations to be used in each image, with data ideally collected from the same equipment; otherwise they may require retraining on additional datasets. ‘Interpretable’ machine learning models or ‘explainable AI’ provide potential solutions to this, but have yet to achieve widespread application^(32,37,38)^.

Segmented and classified multiplex images can be viewed as marked point processes, in which (*x, y*) coordinates representing cell centres are labelled with a ‘mark’ describing their cell type. Statistical and mathematical methods for analysing this data are typically more amenable to interpretation than machine learning approaches, since they quantify interactions between specific cell populations. For example, statistics such as the mean minimum distance between two cell types provide an accessible entry point for analysis of spatial data (e.g., ^(39,40)^), and are available in several software tools^(12,44)^. Statistical approaches based on correlation metrics that were originally developed for ecological applications can also be used to determine whether pairs of cells are colocated more (or less) frequently than would be expected through random chance^(7,45)^. By viewing a multiplex image as a network in which two cell centres are connected if the cells are in physical contact, methods from network science can be used to identify common, recurring motifs within the cell interactions^(7)^. Notably, many network-based approaches use Graph Neural Networks to analyse the spatial patterns formed by the different cell populations (see, e.g., ^(47,48)^). Recently, topological data analysis (TDA), a mathematical field which quantifies the shape of datasets, has emerged as a powerful tool for characterising histology data across multiple scales of resolution in terms of topological features such as connected components and ‘loops’^(46,49)^.

A range of spatial statistics can be used to analyse point processes. These include the Morisita-Horn index, which quantifies dissimilarity between two populations^(40,41)^; Ripley’s K function, which describes clustering or exclusion between points^(15,42)^; and the J-function, which identifies clustering or exclusion by computing nearest-neighbour distributions^(43,50)^. For points with more complex, continuous marks, such as cell size or marker intensity, methods such as mark correlation functions^(16–19)^ or mark variograms^(20,21)^ can be used.

While the above methods have been successfully applied to histology data, the complexity of multiplex imaging data means that there is scope for more detailed statistical and mathematical analyses which surpass what is possible with existing methods. In this paper, we focus on one spatial statistic - the cross-pair correlation function (cross-PCF) - which we use as a foundation to show how existing tools can be adapted to create new statistics that provide more detailed descriptions of multiplex imaging data. The PCF quantifies colocalisation and exclusion between pairs of points, across multiple length scales. It is closely related to the cross-PCF, which identifies correlation between cells of different types. PCF approaches are useful, but their limitations restrict their wider applicability to multiplex data:

1. Cross-PCFs cannot easily resolve heterogeneity in spatial clustering within a region of interest (ROI). Variants of the cross-PCF that account for such heterogeneity do not quantify the contributions of different sub-regions of an ROI to its overall signal^(15)^.
2. Cross-PCFs can identify correlations between pairs of cells in a spatial neighbourhood, but not between 3 or more cell types.
3. Cross-PCFs require cell marks to be discrete, or categorical. Several alternative methods can accommodate continuous marks (e.g., ^(16,20,21)^), but are unsuitable for establishing how the spatial association between cells changes as their continuous marks vary.

In this paper, we discuss three extensions of the cross-PCF that address these limitations. The Topographical Correlation Map (TCM) identifies heterogeneity in the correlation between pairs of cells across an ROI, and has previously been applied by us to imaging mass cytometry data^(7)^. The Neighbourhood Correlation Function (NCF) extends the cross-PCF to quantify correlation between 3 or more different cell types. Finally, the weighted Pair Correlation Function (wPCF) quantifies correlation between two cell populations where one, or both, have a continuous mark, and has been applied to synthetic data^(26)^. In this paper, we present the first applications of the NCF and the wPCF to multiplex imaging data.

The remainder of the paper is structured as follows. In the methods section, we define the TCM, NCF and wPCF, and present motivating examples generated from synthetic data. We also introduce a biological dataset that derives from multiplex IHC images of a murine model of colorectal cancer^(25)^. In the results section, we apply the TCM, NCF and wPCF to this ROI, and demonstrate how each statistic identifies different properties of the spatial interactions that exist between different immune cell populations and cancer cells. We conclude by discussing how these methods expand the scope of the cross-PCF for analysing multiplex images, and suggest possible directions for further investigation.

## Methods

In this section, we introduce the synthetic and experimental datasets which we analyse in this paper. We then define the PCF and cross-PCF, and their extensions: the Topographical Correlation Map (TCM), Neighbourhood Correlation Function (NCF), and weighted Pair Correlation Function (wPCF). The definitions are accompanied by illustrative examples based on the synthetic datasets.

### Data

We construct two synthetic datasets, which are used in the Methods section to develop intuition and understanding of the different spatial statistics. We also introduce a murine colorectal cancer imaging dataset, which is used in the Results section to illustrate the performance of the methods on multiplex imaging data.

### Synthetic data

#### Synthetic dataset I

We consider two cell types, with categorical marks *C*_1_ and *C*_2_. We generate point clouds using different point processes on the left and right hand sides of a 1000 *μ*m × 1000 *μ*m square domain (see Figure 2A and Figure 4A). On the left half of the domain (i.e., for *x* ≤ 500), a Thomas point process is used to generate clustered data^(22)^. This modified Neyman-Scott process samples cluster centres from a Poisson process and samples a fixed number of points from Gaussian distributions around each cluster centre^(23)^. In Synthetic dataset I, we randomly position 20 cluster centres in *x* ≤ 500, and sample 10 points of each cell type from a 2D Gaussian distribution, with standard deviation σ = 20 and mean *μ* located at the cluster centre. In *x* > 500, the same process is used, but 10 cluster centres are chosen independently for each cell type, leading to a composite point pattern containing 300 cells of each type. By construction, synthetic dataset I exhibits strong colocalisation between cells of types *C*_1_ and *C*_2_ in *x* ≤ 500, while each cell type is located in separate clusters in *x* > 500. We assign a second, continuous mark *m* to cells of type *C*_2_. Those with *x* ≤ 500 are randomly assigned a continuous mark *m* ∈ [0, 0.5] while those with *x* > 500 are assigned a mark *m* ∈ (0.5, 1]. Consequently, when a cluster contains both cell types, cells of type *C*_2_ have low marks (*m* ≤ 0.5), and when it contains only cells of type *C*_2_ high marks (*m* ≥ 0.5) are present.

#### Synthetic dataset II

The second synthetic dataset comprises two distinct point patterns, each containing cells of types, *C*_1_, *C*_2_ and *C*_3_ (see Figure 3). In both patterns, three cluster centres are positioned at (*x, y*) = (200, 200), (500, 800), (800, 200). For the first point cloud, each cluster contains 25 cells from two different cell types, with locations chosen from a 2D normal distribution (mean *μ* at the cluster centre, standard deviation σ = 50), so that all three pairwise combinations of cell types are represented (for a total of 50 cells of each type). The same process is used to generate the second point cloud, except all three cell types are present in each cluster (i.e., a total of 75 cells of each type). By contrast, in the first pattern, no cluster contains all three cell types but each pairwise combination of cell types is present in one cluster.

### Multiplex Immunohistochemistry

#### Animals

Intestinal tumour tissue from a villinCre^ER^Kras^G12D/+^Trp53^fl/fl^Rosa26^N1icd/+^ (KPN) mouse was used^(25)^. Procedures were conducted in accordance with Home Office UK regulations and the Animals (Scientific Procedures) Act 1986. Mice were housed individually in ventilated cages, in a specific-pathogen-free (SPF) facility, at the Functional Genetics Facility (Wellcome Centre for Human Genetics, University of Oxford) animal unit. All mice had unrestricted access to food and water, and had not been involved in any previous procedures. The strain used in this study was maintained on C57BL/6J background for ≥ 6 generations.

#### Multiplex immune panel and image pre-processing

Akoya Biosciences OPAL Protocol (Marlborough, MA) was employed for multiplex immunofluorescence staining on FFPE tissue sections of 4-*μ*m thickness. The staining was performed on the Leica BOND RXm auto-stainer (Leica Microsystems, Germany). Six consecutive staining cycles were conducted using primary antibody-Opal fluorophore pairs. The marker panel used is shown in Table 1.

**Table 1:** List of markers and Opals used in the multiplex panel.

| Marker | Opal |
| --- | --- |
| Ly6G (1:300, 551459; BD Pharmingen) | Opal 540 |
| CD4 (1:500, ab183685; Abcam) | Opal 520 |
| CD8 (1:800, 98941; Cell Signaling) | Opal 570 |
| CD68 (1:1200, ab125212; Abcam) | Opal 620 |
| FoxP3 (1:400, 126553; Cell Signaling) | Opal 650 |
| E-cadherin (1:500, 3195; Cell Signaling) | Opal 690 |

The tissue sections were incubated with primary antibody for an hour, and the BOND Polymer Refine Detection System (DS9800, Leica Biosystems, Buffalo Grove, IL) used to detect the antibodies. Epitope Retrieval Solution 1 or 2 was applied to retrieve the antigen for 20 min at 100°C, in accordance with the standard Leica protocol, and, thereafter, each primary antibody was applied. The tissue sections were subsequently treated with spectral DAPI (FP1490, Akoya Biosciences) for 10 minutes and mounted with VECTASHIELD Vibrance Antifade Mounting Medium (H-1700-10; Vector Laboratories) slides. The Vectra Polaris (Akoya Biosciences) was used to obtain whole-slide scans and multispectral images (MSIs). Batch analysis of the MSIs from each case was performed using inForm 2.4.8 software, and the resultant batch-analysed MSIs were combined in HALO (Indica Labs) to create a spectrally unmixed reconstructed whole-tissue image. Cell segmentation and phenotypic density analysis was conducted thereafter across the tissue using HALO.

#### Region of interest overview

We consider a 1mm × 1mm region of interest (ROI) from a KPN mouse intestinal tumour, shown in Figure 1A (three additional regions from this tumour are included in the supporting information). Each colour channel corresponds to a different marker (Blue - DAPI; Orange - CD4; Green - CD68; Magenta - Ly6G; Maroon - FoxP3; Red - CD8; White - E-Cadherin). To obtain a labelled point cloud, individual cell boundaries were identified via cell segmentation (HALO, panel B). Classification of cell types was achieved by considering the average pixel intensity within a cell boundary for each marker individually (e.g., CD4 pixel intensity, panel C), with combinations of cell markers defining different cell types as outlined in Table 2. The final marked point pattern (panel D) was obtained by assigning cell labels to the centroids associated with each cell boundary.

**Table 2:**
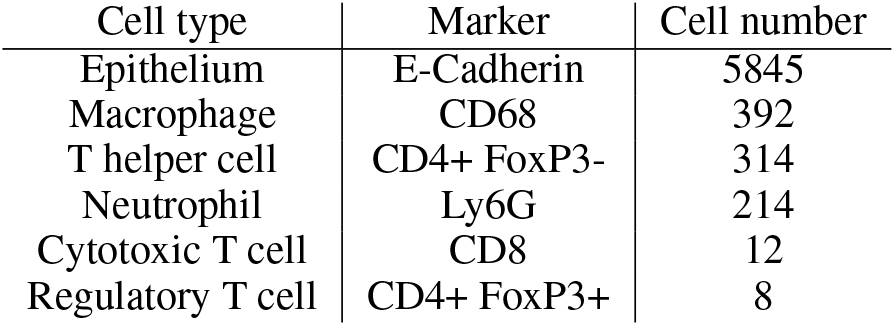
Cell types present in the ROI, with markers and number of cells present. Note that all cells must also contain sufficient DAPI staining to be classified as a cell. Due to low numbers of cytotoxic T cells and regulatory T cells, we exclude them from subsequent analyses.

| Cell type | Marker | Cell number |
| --- | --- | --- |
| Epithelium | E-Cadherin | 5845 |
| Macrophage | CD68 | 392 |
| T helper cell | CD4+ FoxP3- | 314 |
| Neutrophil | Ly6G | 214 |
| Cytotoxic T cell | CD8 | 12 |
| Regulatory T cell | CD4+ FoxP3+ | 8 |

**Figure 1:**
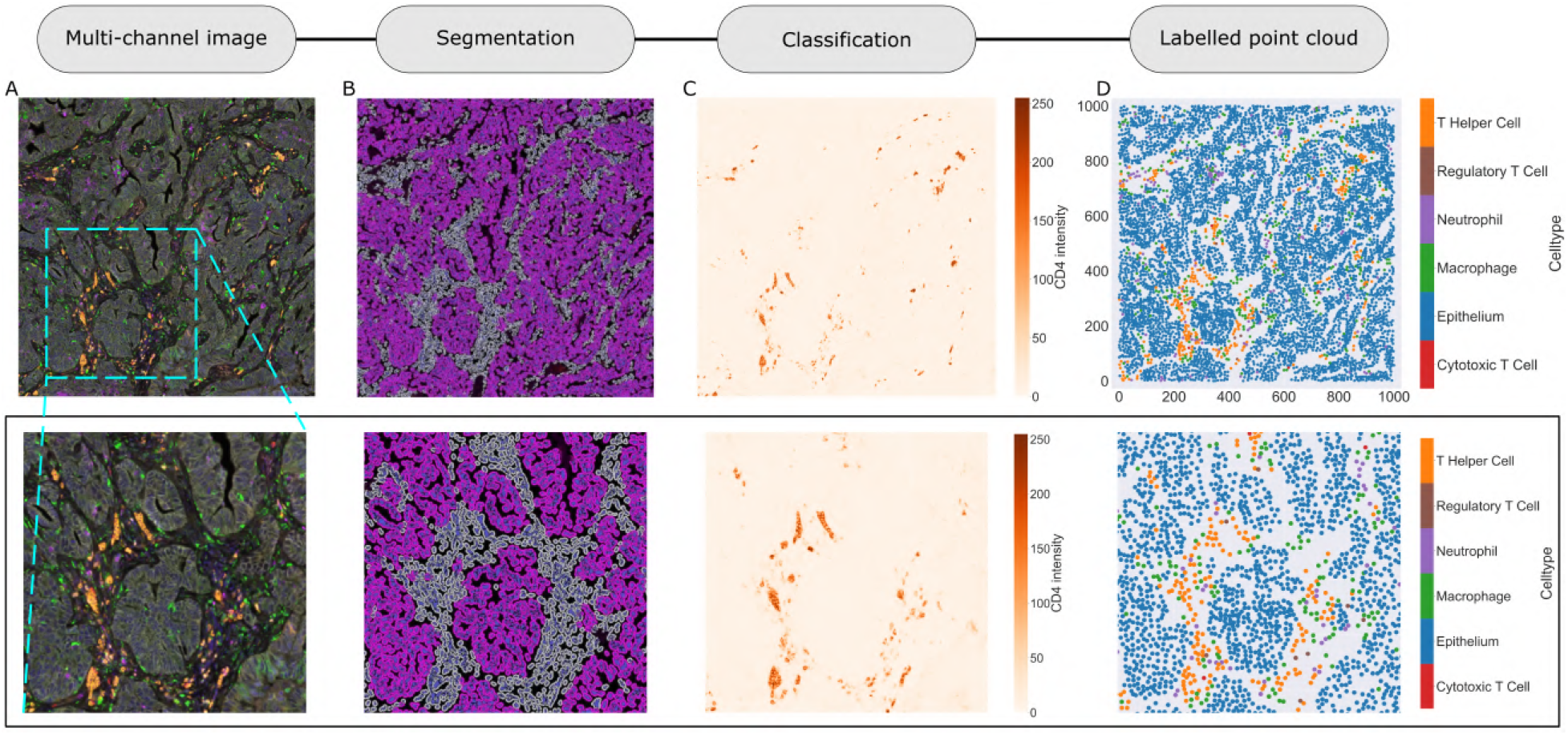
Obtaining point cloud data from a multiplex image. A: 1mm × 1mm ROI from a multiplex immunohistochemistry image of murine colorectal carcinoma (Blue - DAPI; Orange - CD4; Green - CD68; Magenta - Ly6G; Maroon - FoxP3; Red - CD8; White - E-Cadherin). The epithelial cells (E-Cadherin+) are cancer cells which form dense ‘tumour nests’ that are surrounded by stromal regions. Immune cells are largely restricted to the stroma between tumour nests, so the region shows spatial correlation between immune cell subtypes (particularly macrophage, neutrophil and T helper cell) within the stroma, and anti-correlation between immune cells and epithelial cells. B: Cell segmentation (HALO) for the region in panel A. The edges of E-Cadherin positive cells are shown in pink to aid comparison with panel A. C: Pixel intensity from the colour channel corresponding to the CD4 stain only. D: Composite point cloud formed by classifying each cell type stained in panel A, with points placed at the centroids of segmented cells. Lower row: Magnified 500*μ*m × 500*μ*m zoom from the upper panels

**Figure 2:**
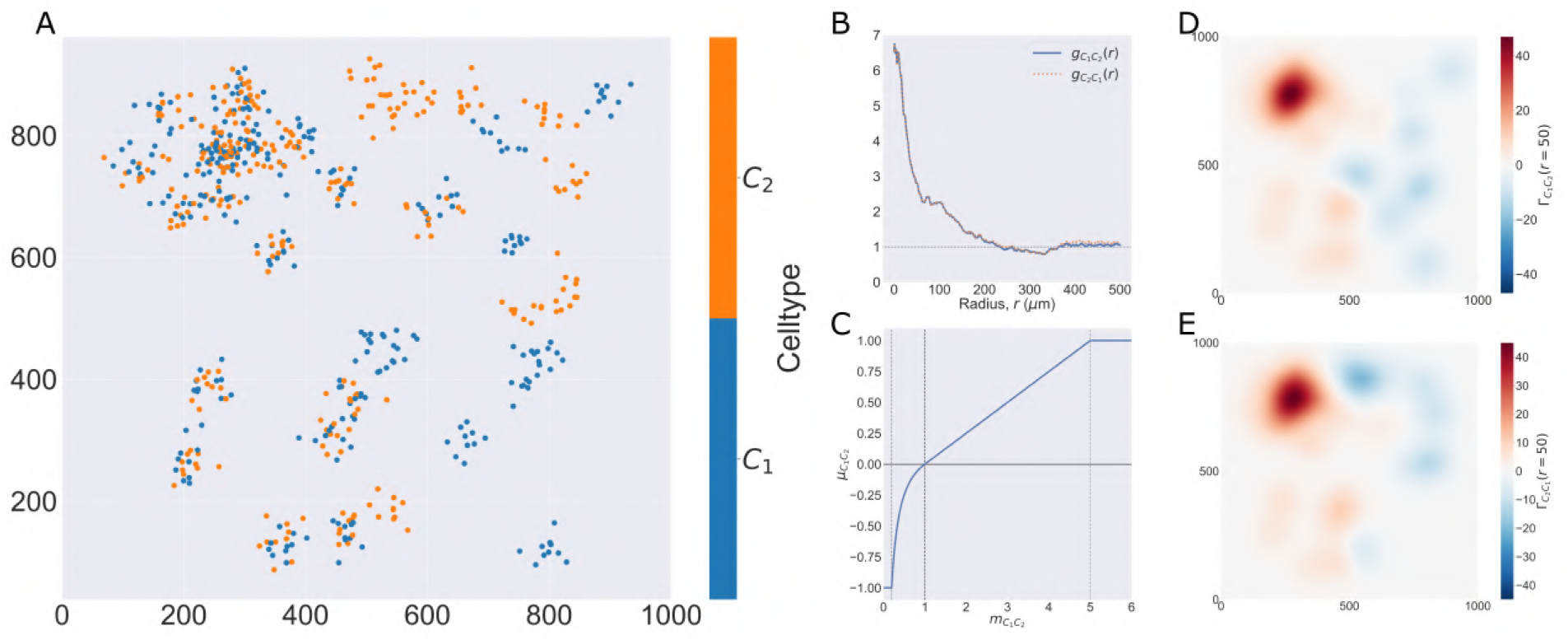
Motivating example I: Cross-PCF and Topographical Correlation Map. A: Synthetic Dataset I: a synthetic point pattern involving 2 cell types, with labels *C*_1_ and *C*_2_. For 0 ≤ *x* ≤ 500, points with labels *C*_1_ and *C*_2_ cluster together; for 500 < *x* ≤1000, points of types *C*_1_ and *C*_2_ form distinct, homogeneous clusters. B: The cross-PCFs 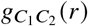 and 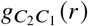 for the point pattern in A. The cross-PCF detects the short range clustering between cells of types *C*_1_ and *C*_2_, which is present for 0 ≤ *x* ≤ 500. The cross-PCFs are almost identical, differing only for large *r* because of boundary correction terms. C: Function used to linearise the mark *m*_*C*1*C*2_ in Equation (6), used to calculate the TCM, for α = 5. Dashed lines represent 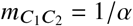, which correspond to the maximum detectable exclusion, CSR, and the maximum detectable clustering. D, E: TCMs 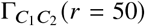 and 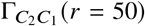. The TCM identifies colocalisation between cells of types *C*_1_ and *C*_2_ in 0 ≤ *x* ≤ 500, and distinguishes between the dense cluster in the top left quadrant and smaller clusters in the bottom left quadrant. The TCM also identifies exclusion between the two cell populations in 500 ≤ *x* ≤ 1000 and shows this to be less pronounced than the clustering in 0 ≤ *x* ≤ 500. Note that while the regions of positive correlation are similar between panels D and E, the regions of negative correlation differ

**Figure 3:**
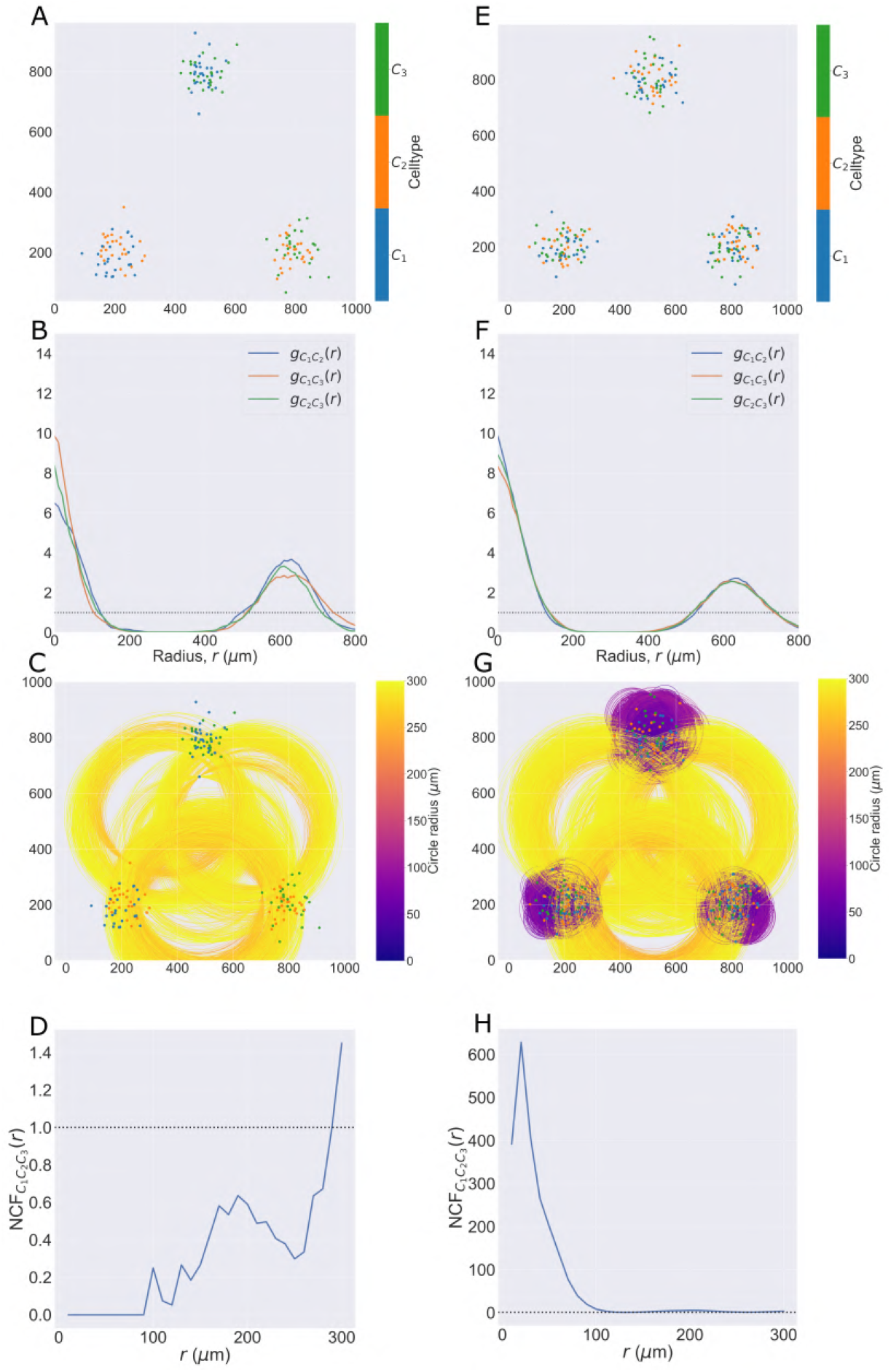
Motivating example II: Neighbourhood Correlation Function. A, E: Synthetic Dataset II: point patterns in which three cell types are spatially correlated pairwise (A) or in triplets (D). In A, each cluster contains only two cell types, do that all three cell types are never in close proximity. In D, all three cell types are in close proximity in each cluster. Hence, in both point patterns there is positive correlation between pairwise combinations of cell types, but the three-way correlations differ between the panels. B, F: Cross-PCFs for the point patterns in panels A and D respectively. These cross-PCFs appear identical, showing strong short-range correlation between the cell types (inside a cluster), exclusion from *r* = 0.2 to *r* = 0.4, and a second peak of correlation around *r* = 0.6 (between clusters). C, G: Minimum enclosing circle for every combination of three points with marks *C*_1_, *C*_2_ and *C*_3_ (up to circles with a radius of *r* = 0.3). Circles with small radii arise when all three cell types are in close proximity (panel G). Circles are coloured according to their radius. D, H: NCFs for the point patterns in A, D respectively. The NCF in panel C correctly identifies short-range exclusion between the three cell types in A, while the NCF in F identifies strong short-range correlation between the three cell types

**Figure 4:**
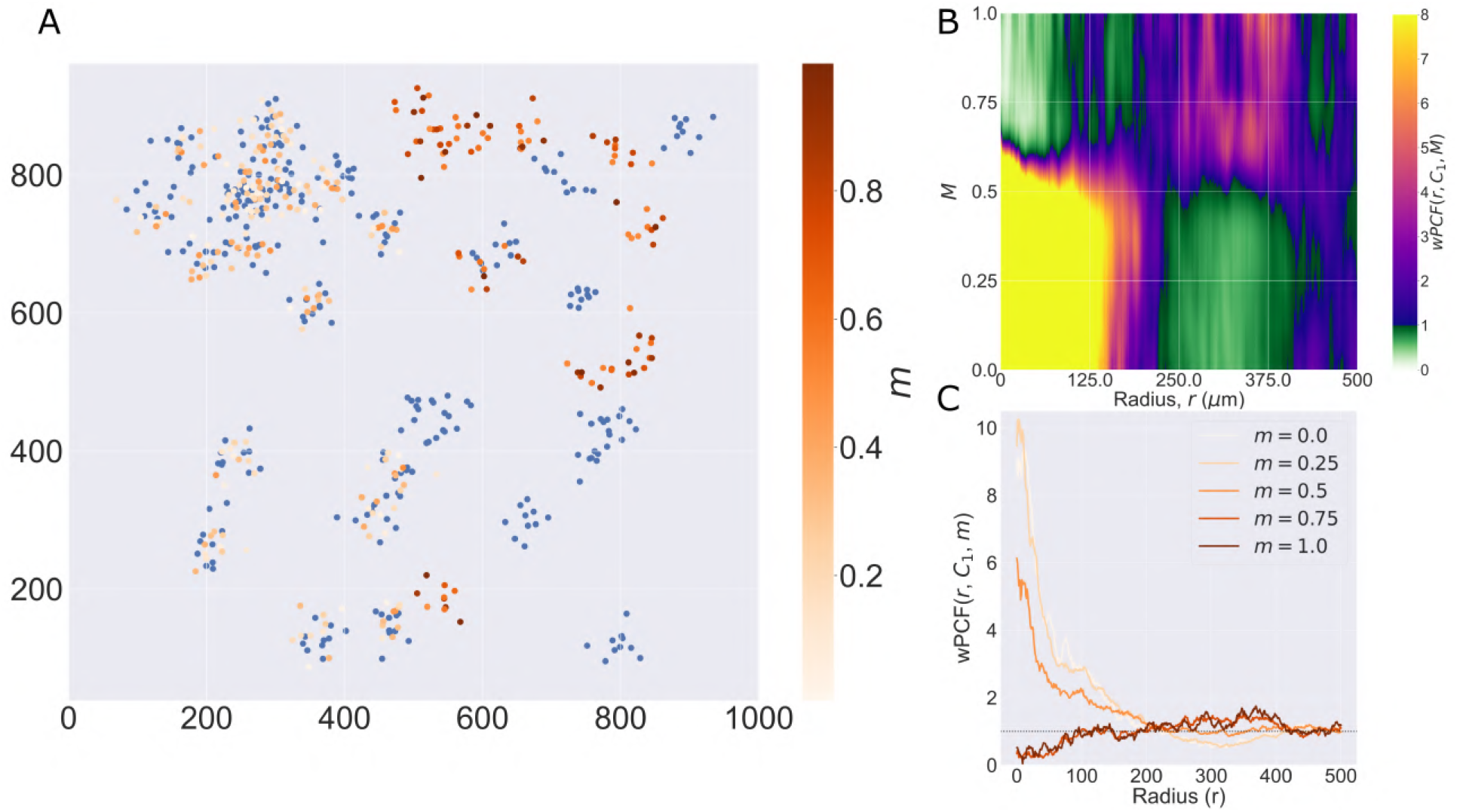
Motivating example III: weighted PCF. A: Synthetic Dataset I: the same point pattern from Figure 2, now shown with the continuous mark *m* associated with cells of type *C*_2_. Recall cells of type *C*_2_ with 0 ≤ *x* ≤ 500 have 0 ≤ *m* <0.5, while those with 500 < *x* ≤1000 have 0.5 ≤ *m* ≤ 1. B: The wPCF, *wPCF* (*r, C*_1_, *m*), for the point pattern in panel A identifies differences in clustering between cells of type *C*_1_ and cells of type *C*_2_ with marks above or below *m* = 0.5. C: Cross-sections of the wPCF in panel B. These plots distinguish the strong clustering of cells of type *C*_1_with cells of type *C*_2_ that have *m* < 0.5 and their weak exclusion from cells of type *C*_2_ that have *m* > 0.5.

The ROI in Figure 1 was selected because of the clear separation between the spatial position of immune cell subtypes and tumour nests (epithelial cells), with immune cells located predominantly in regions between epithelial cell islands. Table 2 summarises the different cell types, the markers used to define them, and the number of cells of each type in the ROI.

### Spatial statistics

We consider a point pattern in a rectangular domain Ω = [0, 1000*μ*m] × [0, 1000*μ*m]. We note that, with suitable modifications, the statistical analysis can be applied to more complex regions (e.g., with holes) and higher dimensional domains. The point pattern comprises *N* points (or cells). Cell *i* (*i* ∈ 1, 2, …, *N*) has spatial location **x**_**i**_ = (*x*_*i*_, *y*_*i*_), and a set of marks which may be categorical (e.g., a label for a cell type, or a true/false label indicating whether a cell’s average stain intensity exceeds a threshold value), or continuous (e.g., the average stain intensity of a particular mark within a cell). For clarity, we denote categorical and continuous marks by *c* and *m* respectively. We use lowercase for marks associated with a particular point and uppercase for target values.

We introduce the indicator function 𝕀(*C, c*) to determine whether a categorical mark associated with a point matches a target mark:

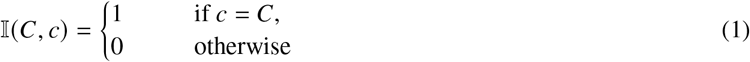

When we define correlation functions below, we will need to determine whether two points are separated by a distance ‘close to’ *r*. Accordingly, we introduce the following function, *I*_*k*_ (*r*):

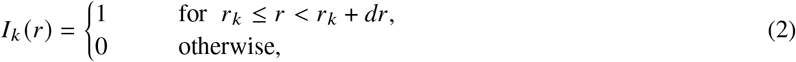

where *r*_*k*_ is the inner radius of an annulus of width *dr*. We consider a sequence of annuli, with *r*_*k*+1_ = *r*_*k*_ + *dr, dr* > 0 and *r*_0_ = 0. We denote by *A*_*k*_ (**x**) the area of the annulus with inner radius *r*_*k*_ centred at the point **x**. If this annulus lies wholly inside the domain then 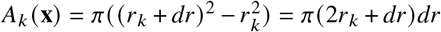; otherwise, only the area contained within the domain is recorded.

### Pair Correlation Function (PCF)

#### Aims

The Pair Correlation Function (PCF), *g*(*r*), quantifies spatial clustering or exclusion between pairs of points separated by a distance *r* within an ROI, compared to a suitably selected null distribution. While a range of null distributions could be considered (e.g., using a Matérn hard core process to simulate randomly distributed cell centres separated by a minimum distance to approximate a cell radius^(24)^), we assume the null distribution is complete spatial randomness (CSR) as represented by a homogeneous spatial Poisson point process with intensity *λ* > 0 chosen to match the intensity of the point pattern being analysed.

**Definition** Let 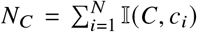 be the number of points in Ω with *c*_*i*_ = *C*, for some categorical mark *C*. The PCF, *g*(*r*), is defined as follows:

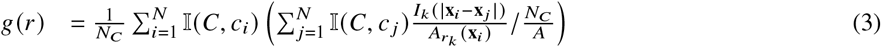

where *A* is the total area of the domain Ω. While there are many ways to account for edge effects associated with points close to the domain boundary, we account for them by adjusting the contribution of each point to account for the area of annulus *k* contained within the domain, 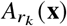.

We note from Equation (3) that *g*(*r*) = 1 corresponds to CSR. Further, if *g* (*r*) > 1 then points separated by distance *r* are observed more frequently than expected under CSR and we say that points at this lengthscale are clustered relative to CSR. Similarly, *g*(*r*) < 1 indicates fewer points than expected and is interpreted as exclusion at lengthscale *r*.

The structure of Equation (3) provides the basis for the generalisations of the PCF introduced below.

### Cross Pair Correlation Function (Cross-PCF)

#### Aims

The cross-PCF describes correlation between pairs of points separated by distance *r* which may have different categorical labels.

#### Definition

Consider the categorical marks *C*_1_ and *C*_2_. The cross-PCF, 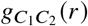, is defined as follows:

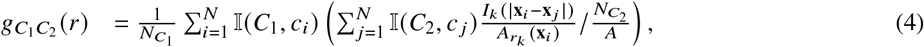

where 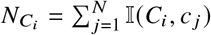 is the number of points with mark *C*_*i*_. We note that when *C*_1_ = *C*_2_, Equation (4) reduces to Equation (3) (i.e., the cross-PCF reduces to the PCF).

#### Example

The interpretation of the cross-PCF is similar to that for the PCF, with 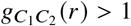 1 indicating correlation between points with marks *C*_1_ and *C*_2_ separated by distance *r* and 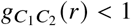 indicating exclusion at distance *r*.

In Figure 2, we compute two cross-PCFs for Synthetic Dataset I. In Figure 2A, cells with labels *C*_1_ and *C*_2_ are strongly spatially correlated on the left half of the domain, while they are clustered separately on the right half. Figure 2B shows the cross-PCFs 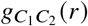 and 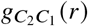 for this point pattern. Colocalisation between the cell types is identified for *r* ⪅ 200. The cross-PCFs are almost identical, since the cross-PCF is symmetric up to boundary correction terms. While the cross-PCF successfully identifies the presence of clustering between the two cell types, it does not provide information about differences in colocalisation on the left and right hand sides of the domain.

### Topographical Correlation Map (TCM)

#### Aims

The Topographical Correlation Map (TCM), 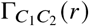, was introduced by us in^(7)^ to visualise spatial heterogeneity in the correlation between pairs of points across an ROI. Motivated by Equation (4), each point with mark *C*_1_ is assigned a value that quantifies its correlation with points with mark *C*_2_. A series of kernels centred at each point with mark *C*_1_ is summed to produce a spatial map of local correlations between the cell types. We note that, since these kernels are centred on points marked *C*_1_, the TCM is not symmetric (i.e., 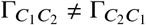 if *C*_1_ ≠ *C*_2_).

#### Definition

The TCM, 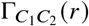, is visualised at a specific length scale *r*. Unless stated otherwise, we fix *r* = 50*μ*m which corresponds to clustering on the length scale of 2-3 cell diameters. We associate a continuous mark 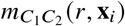 with each cell *i* with mark *C*_1_, such that

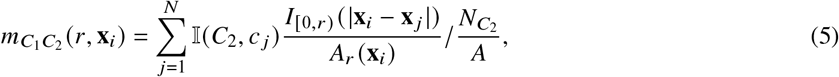

where *A*_*r*_(**x**_*i*_) is the area of that part of the circle with radius *r μ*m centred at **x**_*i*_ that falls within the ROI. 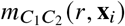 can be viewed as the contribution of each point *i* to the cross-PCF, 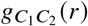, for the special case of an annulus with inner radius 0 and width *dr* = *r*. Thus 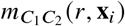 is interpreted similarly to the cross-PCF: 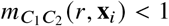 indicates anti-correlation between cells with marks *C*_1_ and *C*_2_ separated by a distance of at most *r μ*m, and 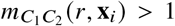 indicates correlation.

Since 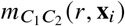 is based on a ratio of observed counts to counts expected under CSR, its interpretation is nonlinear: an observation of three times as many points as expected corresponds to 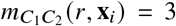, while three times fewer points than expected leads to 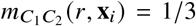. To facilitate interpretation, we rescale 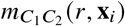 to produce a transformed mark 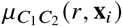 in which clustering and exclusion can be compared on a linear scale, with 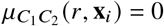 when 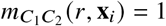:

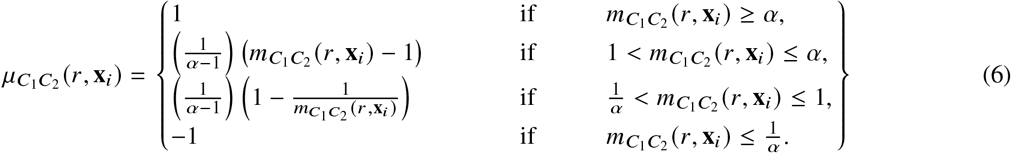

In Equation (6), the constant α > 1 describes the maximal degree of clustering (or exclusion) which can be resolved under this transformation. A sketch of Equation (6) is presented in Figure 2C, for α = 5 (henceforth, we fix α = 5).

After calculating 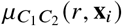 for each cell with mark *C*_1_, we centre a Gaussian kernel with standard deviation σ = *r μ*m and maximum height 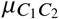 at **x**_*i*_. The TCM, 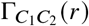, is obtained by summing over all cells with mark *C*_1_ in the domain:

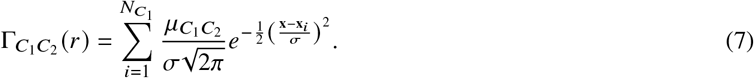

Regions in which the TCM is positive indicate that more points marked *C*_1_ are positively correlated with points marked *C*_2_ in this area than would be expected under CSR, at lengthscales up to *rμ*m. Similarly, the TCM is negative in regions where points with mark *C*_1_ are negatively correlated with points with mark *C*_2_.

#### Example

Figure 2D and E show TCMs associated with Synthetic Dataset I (the point pattern in Figure 2A) for *r* = 50. Panel D shows 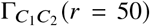 and panel E shows 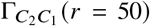. Both TCMs identify differences in the colocalisation of the two cell types on the left and right sides of the domain. In particular, 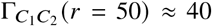 in the upper left quadrant of panels D and E, indicating strong positive correlation, with weak association in the lower left (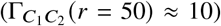). For *x* ≥ 500 both TCMs correctly identify weak anti-correlation (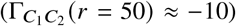) between cells of types *C*_1_ and *C*_2_. The cross-PCFs in panel B are dominated by the correlation on the left hand side of the domain, and are unable to resolve the heterogeneity in clustering between the left and right sides of the domain. We note that 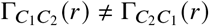, since the kernels used to construct the TCM are centred on cells with label *C*_1_ (and vice versa). While areas in which cells with mark *C*_1_ and mark *C*_2_ are co-located are identified by positive values of both 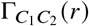 and 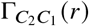, their values differ in regions where one or other TCM is negative, as in these regions the cell densities vary (e.g., on the right hand side of panels D and E). We, therefore, emphasise that 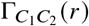 provides a spatial map of subregions in which cells with mark *C*_1_ are correlated (or anti-correlated) with cells with mark *C*_2_. Finally, we note that the TCM is not a density map showing the presence (or absence) of the cell types individually; for example, when 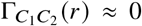, either cells of type *C*_1_ are absent, or cells of both types are present in numbers consistent with CSR.

### Neighbourhood Correlation Function (NCF)

#### Aims

The neighbourhood correlation function (NCF(*r*)) extends the PCF to quantify spatial colocation between 3 or more cell types with different categorical marks. We compare the observed number of triplets of points with marks *C*_1_, *C*_2_ and *C*_3_ within a neighbourhood of size *r* against the number of triplets expected under CSR. Selecting an appropriate definition for such a neighbourhood is non-trivial: while it is straightforward to calculate the Euclidean distance between 2 points, many metrics can be used to calculate the proximity of 3 or more points. We require a metric which is interpretable and extends naturally to more than 3 points. Metrics such as the area of the polygon spanning the points are unsuitable (the area of the polygon is identically zero when all points fall on a straight line, even though the points could be far apart). We consider the minimum enclosing circle (details below) as it requires all cells to lie within a ‘neighbourhood’ of each other (with the distance between any two points at most 2*r*, where *r* is the radius of the minimum enclosing circle).

#### Definition

Consider a point pattern for which there are 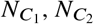, and 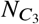 points with categorical marks *C*_1_, *C*_2_ and *C*_3_ respectively. We say that three points from this pattern fall within a ‘neighbourhood’ of radius *r* if there is a circle of radius *r* which encloses all three points. For a given set of three points ζ = {**x**_1_, **x**_2_, **x**_3_}, let *R* (ζ) be the radius of the smallest circle enclosing every point in ζ (the ‘minimum enclosing circle’).

There are 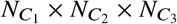 possible triplets containing one point with each mark. We calculate *R* for each of these, and then determine the number of circles of radius *r* containing a unique grouping of cells with each mark (as for the PCF, these values are grouped into discrete bins of width *dr*).

As for the PCF, we compare the number of minimum enclosing circles with radius *r* with the number expected under CSR. The probability of three points lying within a neighbourhood of radius *r, p*_3_(*r*), is:

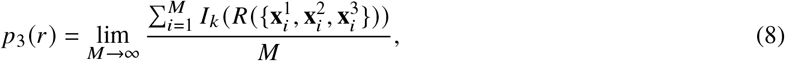

where 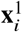, 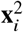 and 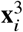 are three points within the domain, sampled under CSR. Since sampling such points is computationally cheap, *p*_3_ *r* can practically be approximated for an arbitrary domain by sampling a large number of random triplets (in this section, we use *M* = 10^7^) and calculating their minimum enclosing circles.

For a point pattern containing *N* points, the NCF is defined as the ratio of the observed number of smallest neighbourhoods of radius *r* to the number of such neighbourhoods expected under CSR, *N*_1_ × *N*_2_ × *N*_3_ × *p*_3_(*r*):

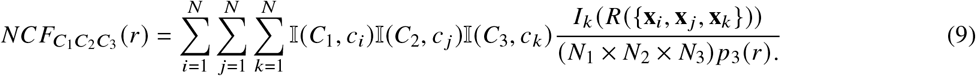

We note that it is straightforward to extend the NCF for *n* categorical marks:

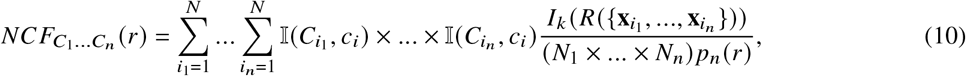

where *p*_*n*_ (*r*) is the probability that *n* points sampled under CSR fall within a minimum enclosing circle of radius *r*.

#### Example

As for the PCF, *NCF* (*r*) > 1 indicates clustering and *NCF* (*r*) < 1 indicates exclusion. We interpret the length scale *r* associated with the NCF as the neighbourhood radius within which the points are contained.

In Figure 3 we compare the cross-PCFs and 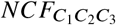 for the two point patterns from Synthetic Dataset II. In Figure 3A, each cluster consists of only two cell types, so that any pairwise combination of cell types can be found in close proximity while all three cell types are never in close proximity. In Figure 3E each cluster contains all three cell types.

Figures 3B and F show that, for Synthetic Dataset II, all cross-PCFs 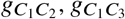 and 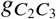 have the same shape and, hence, that pairwise correlation is insufficient to distinguish the two point patterns in this dataset. Figures 3C and G show all minimum enclosing circles (with radius up to 300 *μ*m) for these point patterns, coloured according to the radius of the circle; note that when all three cell types are present within the same cluster, there are a large number of small (purple) circles present. Figure 3D shows that, in this point pattern, all three cell types are never observed within a circle of radius *r* < 100*μ*m. The peak at *r* ≈ 200*μ*m indicates that approximately as many triplets of cells of type *C*_1_, *C*_2_ and *C*_3_ can be enclosed by a circle of radius 200*μ*m as expected under CSR; this corresponds approximately with the distance between the edges of neighbouring clusters. In contrast, Figure 3H shows that the NCF distinguishes the two point patterns, by identifying strong correlation between the three cell types in neighbourhoods with radii of at most 100*μ*m for the three-way correlation point pattern, which corresponds to the approximate radius of the clusters.

### Weighted Pair Correlation Function (wPCF)

#### Aims

The weighted Pair Correlation Function (wPCF) extends the cross-PCF to describe correlation and exclusion between cells marked with labels that may be categorical or continuous. Here, we focus on pairwise comparisons between points marked with a categorical label (e.g., points of type *C*_1_) and those marked with a continuous label (e.g., points with mark *m* ∈ [0, 1]. The wPCF can also compute correlations between points labelled with two continuous marks (see^(26)^ for an example of this).

#### Definition

Consider a set of points labelled with categorical marks (*C*_1_) and continuous marks (*m* ∈ [*a, b*] for some *a, b* ∈ ℝ). The wPCF describes the correlation between points with a given target mark *M* ∈ [*a, b*] and those with a categorical mark *C*_1_, at a range of lengthscales *r*.

The cross-PCF cannot be calculated for such points since, for a continuous mark, 𝕀 (*M, m*) is zero almost everywhere. As such, we replace 𝕀(*C, c*) with a generalised version, the ‘weighting function’ *w*_*m*_ (*M, m*), to account for values of continuous marks *m* that are ‘close to’ a target mark *M* in the following way:

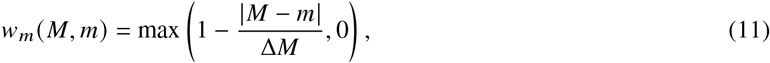

where the positive parameter Δ*M* determines the width of the function’s support. Many other functional forms could be used, provided that they have compact support, *w*_*m*_(*M, M)* = 1 and *w*_*m*_(*M, m*) ∈ [0, 1] (see^(26)^ for details of alternative functional forms and a detailed analysis of how the choice of weighting function influences the signal to noise ratio in the resulting wPCF).

The wPCF is defined as follows:

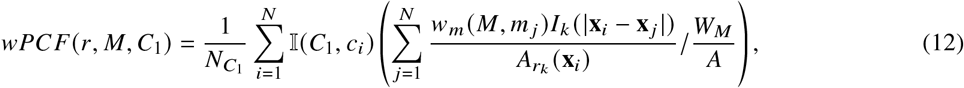

where 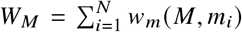 is the total ‘weight’ associated with the target label *M* across all points. The wPCF extends the cross-PCF by weighting the contribution of each point based on how closely its continuous mark matches the target mark.

We note that the wPCF can be used to compare point clouds with two continuous marks by replacing the categorical target mark *C*_1_ with a second continuous mark (say, *M*_1_):

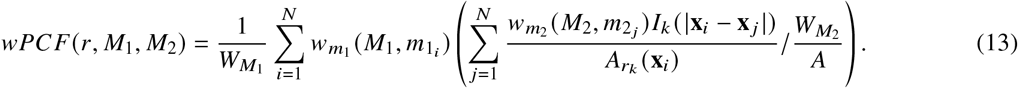

In Equation (13) the weighting functions 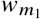 and 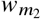 quantify proximity to target marks *M*_1_ and *M*_2_ respectively. Note that since the ranges of the marks *m*_1_ and *m*_2_ may differ substantially, the functions 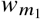 and 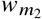 may not necessarily use the same value of Δ*M* in Equation (11) (for an example, see^(26)^).

#### Example

We again use Synthetic Dataset I, where points with *m* < 0.5 are on the left hand side of the domain, and have been placed in clusters with the *C*_1_ cells. In contrast, points on the right hand side have *m* > 0.5 and cluster independently from the *C*_1_ clusters.

Figure 4B shows *wPCF* (*r, C*_1_, *M*) for the point pattern in Figure 4A, with cross-sections of the wPCF shown in Figure 4C. For a given target value *M*, the cross-sections of the wPCF can be interpreted in the same manner as the cross-PCF or PCF. Figure 4B identifies two types of correlation in the data, each associated with different values of *m*. For 0 < *r* ≈ 150, there is strong short-range clustering between cells of type *C*_1_ and cells of type *C*_2_ with *m* < 0.5, with weak short-range exclusion up to this length scale for *m* > 0.5. Since cells on left hand side of the domain have 0 ≤ *m* < 0.5, and those on the right hand side have 0.5 ≤ *m* ≤ 1, this effect is consistent with the information from the cross-PCF and TCM above. One advantage of visualising the wPCF as a heatmap (Figure 4B) is that it identifies threshold values of *M* at which the nature of the cell-cell correlations changes, as demonstrated in^(26)^.

## Results

In this section, we illustrate the utility of the TCM, NCF and wPCF through their application to an ROI from a multiplex immunohistochemistry image of a murine colorectal carcinoma (see methods for details, and Section S1 of the supplementary information for similar analyses of three additional ROIs). Figure 5 shows the cross-PCFs which describe pairwise correlations between all cell types present in the data. Due to the low numbers of cytotoxic and regulatory T cells, we focus subsequent analyses on relationships between epithelium and T helper cells (two abundant cell types which are spatially anti-correlated) and on T helper cells and macrophages (the most abundant immune cell subtypes, which are spatially correlated). When applying the NCF, we include neutrophils as a third immune cell subtype which colocalises with T helper cells and macrophages.

**Figure 5:**
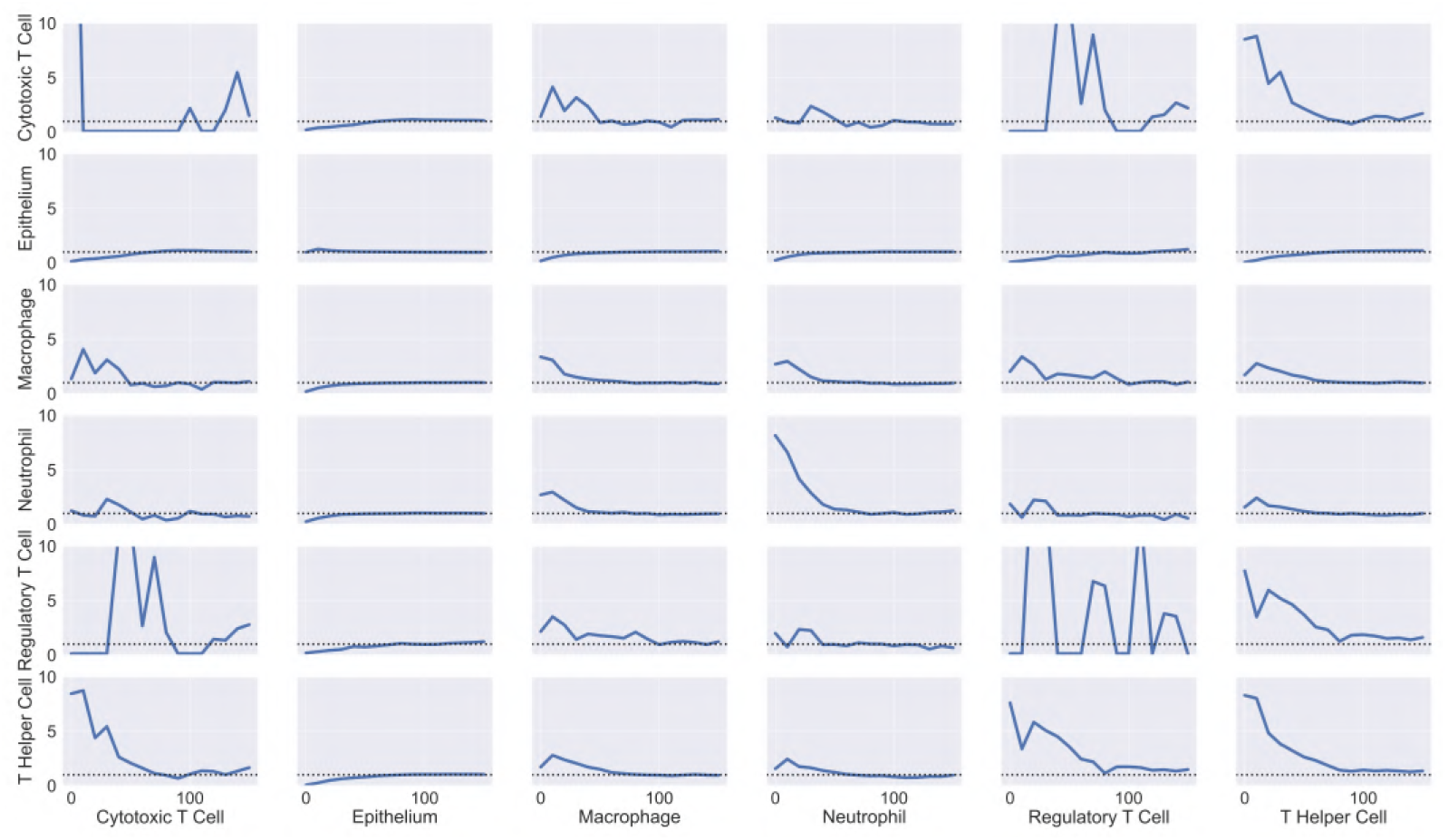
Cross-PCFs for all pairwise combinations of cell types in the ROI. Cross-PCFs for all pairs of cell types from the ROI. We observe exclusion between epithelium and all immune cell subtypes, and strong pairwise correlation with macrophages, neutrophils, and T Helper Cells on short lengthscales. Results involving regulatory and cytotoxic T Cells are noisy as their cell counts are low in this ROI

### Cross-PCFs and TCMs identify colocalisation and exclusion in cell centre data

We first consider T helper cells (Th) and macrophages (M), which are shown to be colocalised from the cross-PCFs in Figure 5. Figure 6A shows the channels of the multiplex image that correspond approximately with T helper cells (CD4+, orange) and macrophages (CD68+, green); the cell centres of these cell populations within the ROI are shown in Figure 6B (see Figure 1 for other cell locations). In this ROI, both T helper cells and macrophages are predominantly found in the stromal tissue between islands of (cancerous) epithelial cells, leading to positive spatial correlation on short length scales (0 ⪅ *r* ⪅ 75*μm*).

**Figure 6:**
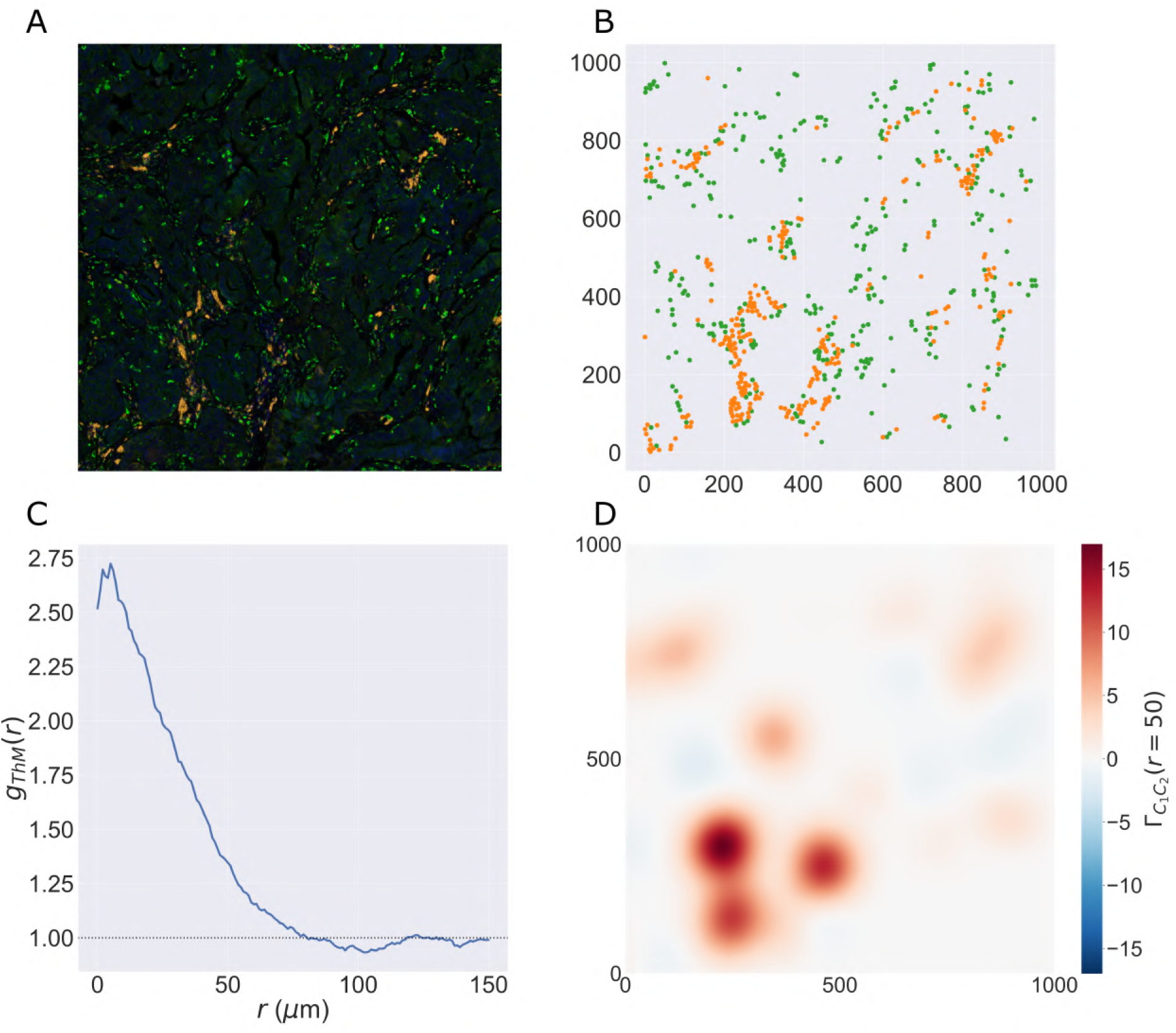
PCF and TCM for positively correlated cell types. A: Locations of T helper cells (CD4+, orange) and macrophages (CD68+, green) in the ROI (with DAPI, blue). These cell types colocalise in the tissue between epithelial cell islands. B: Cell centres identified as T helper cells (orange) and macrophages (green). C: PCF for T helper cells to macrophages, *g*_*ThM*_(*r*). These cell types are spatially colocated over a wide range of distances, i.e., *g*_*ThM*_(*r*) > 1 for 0 ⪅ *r* ⪅ 75*μ*m. D: TCM for T helper cells to macrophages, Γ_*ThM*_, for *r* = 50*μ*m. Red regions indicate colocalisation of the cell types in stromal regions, while blue regions correspond to isolated T helper cells

Colocalisation is clearly identified by the cross-PCF in Figure 6C: for 0 ≤ *r* ⪅ 75, *g*_*ThM*_ (*r*) > 1, indicating clustering between the cells of up to 2.75 times greater than expected under CSR, on length scales up to approximately 75 *μ*m (a distance approximating the width of the stromal region that separates epithelial clusters).

Figure 6D shows Γ_*ThM*_ (*r*) for *r* = 50*μ*m. This permits the clustering identified by the cross-PCF to be mapped onto the ROI, revealing subregions in which T helper cells are spatially colocated with, or excluded from, macrophages. We observe strong clustering in stromal regions, with islands of weak exclusion where isolated T helper cells are present. We conclude that, while T helper cells typically colocalise with macrophages, certain subregions of the ROI that contain T helper cells have low numbers of macrophages within a 50 *μ*m radius. Further, these subregions do not contribute significantly to the overall correlation of T helper cells with macrophages in the cross-PCF.

In Figure 7 we focus on T helper (Th) and epithelial cells (E), which are shown to be anti-correlated by the cross-PCFs in Figure 5. In Figure 7C, the cross-PCF *g*_*ThE*_(*r*) shows exclusion for 0 ≤ *r* ⪅ 75, with the strongest exclusion occurring on small length scales. This exclusion is also identified by Γ_*ThE*_(*r*) in Figure 7D, which shows that the cross-PCF is dominated by strong exclusion from T helper cells in the stromal islands between epithelial cells as expected. The contributions from T helper cells outside the stromal regions (e.g., those in the lower right quadrant of the ROI) are negligible compared to those in the lower left quadrant, due to the large number of T helper cells in that subregion.

**Figure 7:**
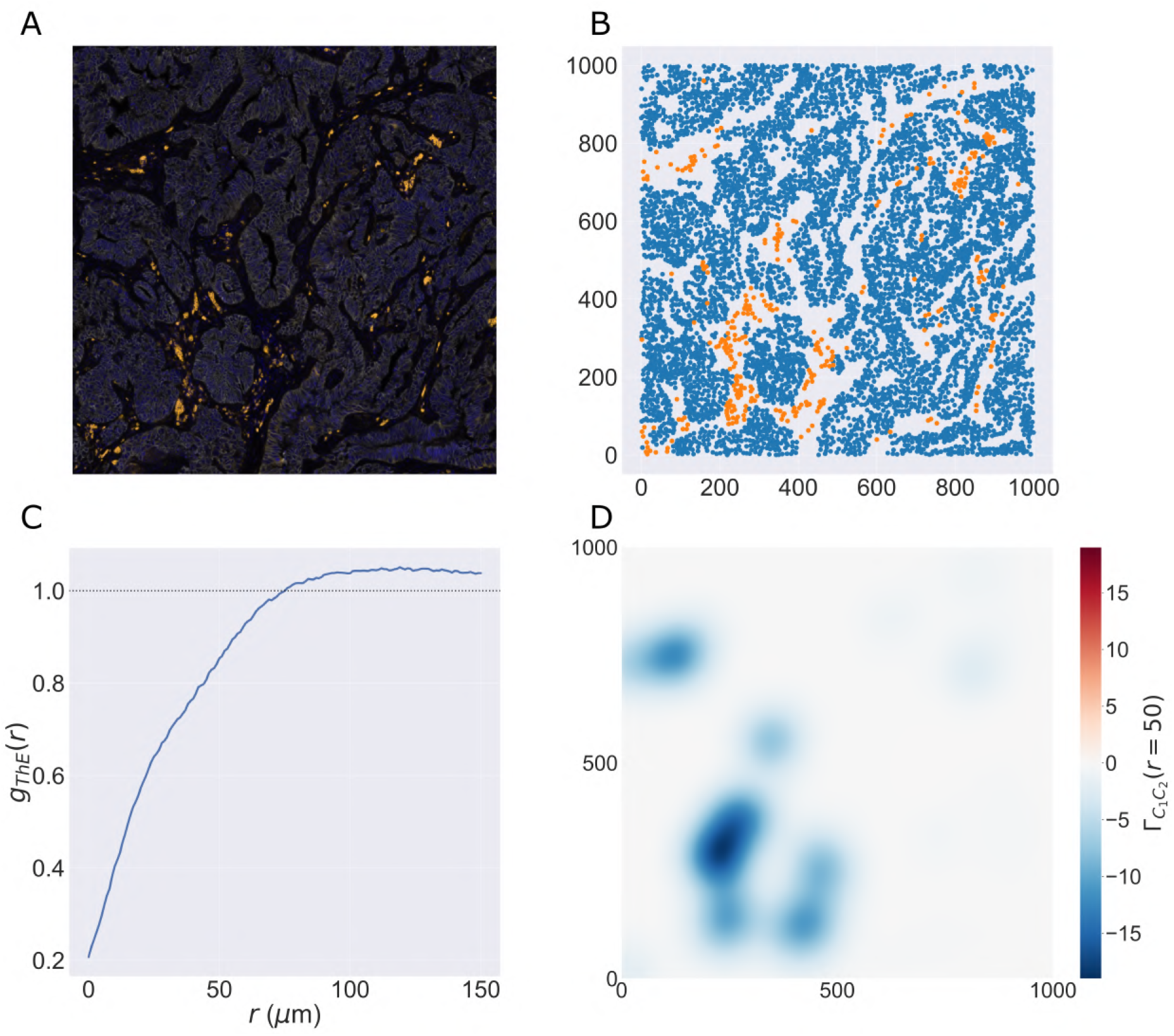
PCF and TCM for negatively correlated cell types. A: Locations of T helper (CD4+, orange) and epithelial cells (E-Cadherin+, white) in the ROI (with DAPI, blue). Epithelial cells exist in clumped ‘nests’, with T helper cells restricted to the stromal regions between them. B: Cell centres of T helper cells (orange) and epithelial cells (blue). C: PCF for T helper cells to epithelial cells, *g*_*ThE*_(*r*). We observe strong spatial exclusion, as *g*_*ThE*_(*r*) < 1 for *r* ⪅ 75. D: TCM for T helper cells to epithelial cells, Γ_*ThE*_(*r*), for *r* = 50*μ*m. The blue regions showing strong exclusion indicate subregions of the ROI which are devoid of epithelial cells. The strongest signals occur where T helper cells are organised in large clusters, while regions with few T helper cells do not contribute significantly to the cross-PCF

### Neighbourhood Correlation Functions identify spatial correlations between 3 cell types simultaneously

Figure 8 shows the NCF for macrophages, T helper cells, and neutrophils (spatial locations shown in Figure 8A). We calculate the smallest circles enclosing each triplet containing one of each cell type, and note their radii. Figure 8B compares the number of circles with radius *r* observed in the data, with the expected number if macrophages, neutrophils and T helper cells are randomly distributed (obtained via bootstrapping as described in the methods, for *M* = 10^8^). More circles are observed than expected under CSR. By taking the ratio of the curves in Figure 8B we generate the NCF in Figure 8C. The NCF shows that triplets comprising a macrophage, a neutrophil and a T helper cell are up to 35 times more likely to cluster within a neighbourhood of radius 0-20*μ*m than would be expected if the cells were randomly distributed. We conclude that these cell types are frequently found together.

**Figure 8:**
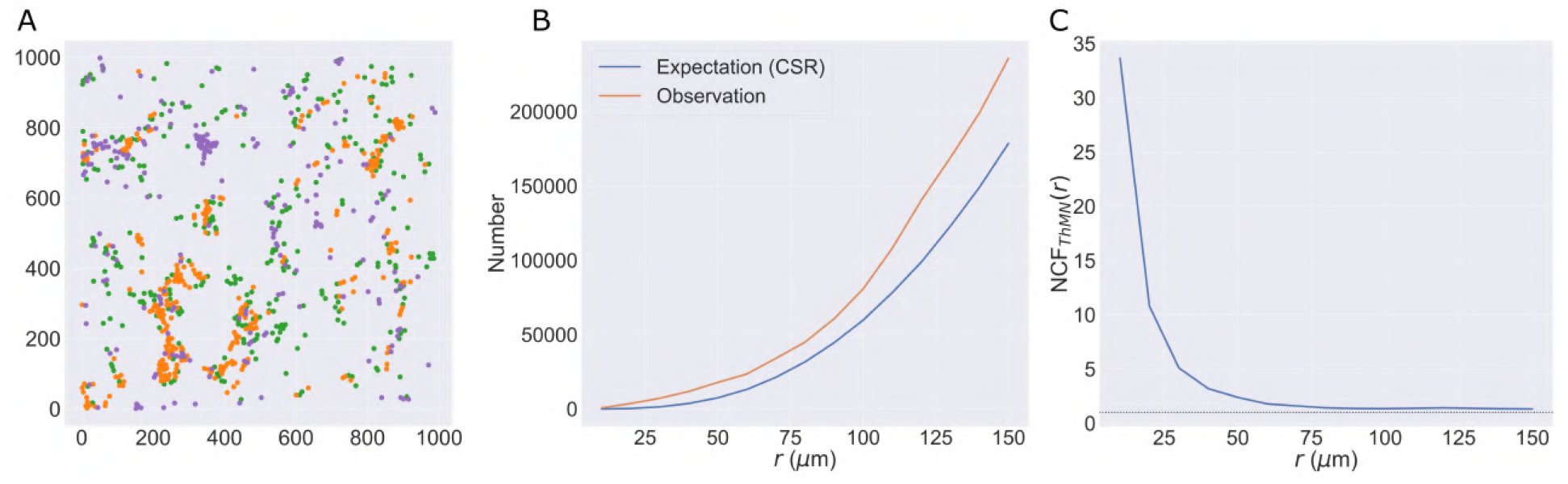
The NCF identifies spatial colocalisation between 3 cell types. A: Locations of T helper cells (orange), macrophages (green) and neutrophils (purple) extracted from the ROI. All three cell types are found in stromal regions, while macrophages and neutrophils are more likely to be observed within the epithelial islands (e.g., in the top left corner). B: Expected and observed numbers of circles of radius *r*. C: NCF obtained by computing the ratio of the curves in panel C, NCF_*ThMN*_(*r*). For *r* ⪅ 75*μ*m, neutrophils, macrophages and T helper cells are colocalised within a circle of radius *r* more often than would be expected under CSR.

### The Weighted Pair Correlation Function identifies correlations without classification or segmentation

Recall that in order to apply the PCF, cross-PCF, TCM, and NCF, the multiplex imaging data must be segmented and then classified to identify cell centres and assign them categorical labels (or cell types). We now show how the wPCF can be applied directly to multiplex imaging data to identify spatial correlations, without segmentation or classification.

Figure 9 demonstrates that the wPCF can identify correlations when some cells are not classified. Rather than specifying a threshold value of the CD4 marker intensity to identify CD4+ cells, we instead view the average CD4 intensity of each cell as a continuous mark. Figure 9A shows epithelial cells determined by specifying a threshold, while Figure 9B shows all cells labelled according to their average CD4 intensity. The wPCF calculated in panel C shows that the spatial positions of cells with low CD4 intensity differ from those with high CD4 expression (with Δ*M* = 2 in Equation (11)). Figures 9C and D show that cells with mean CD4 intensity below approximately 4 are not strongly correlated with epithelial cells. However, for larger values of CD4 intensity, the profiles of the wPCF are in good agreement with the cross-PCF *g*_*ThE*_(*r*) (shown as a red dashed line in Figure 9D).

**Figure 9:**
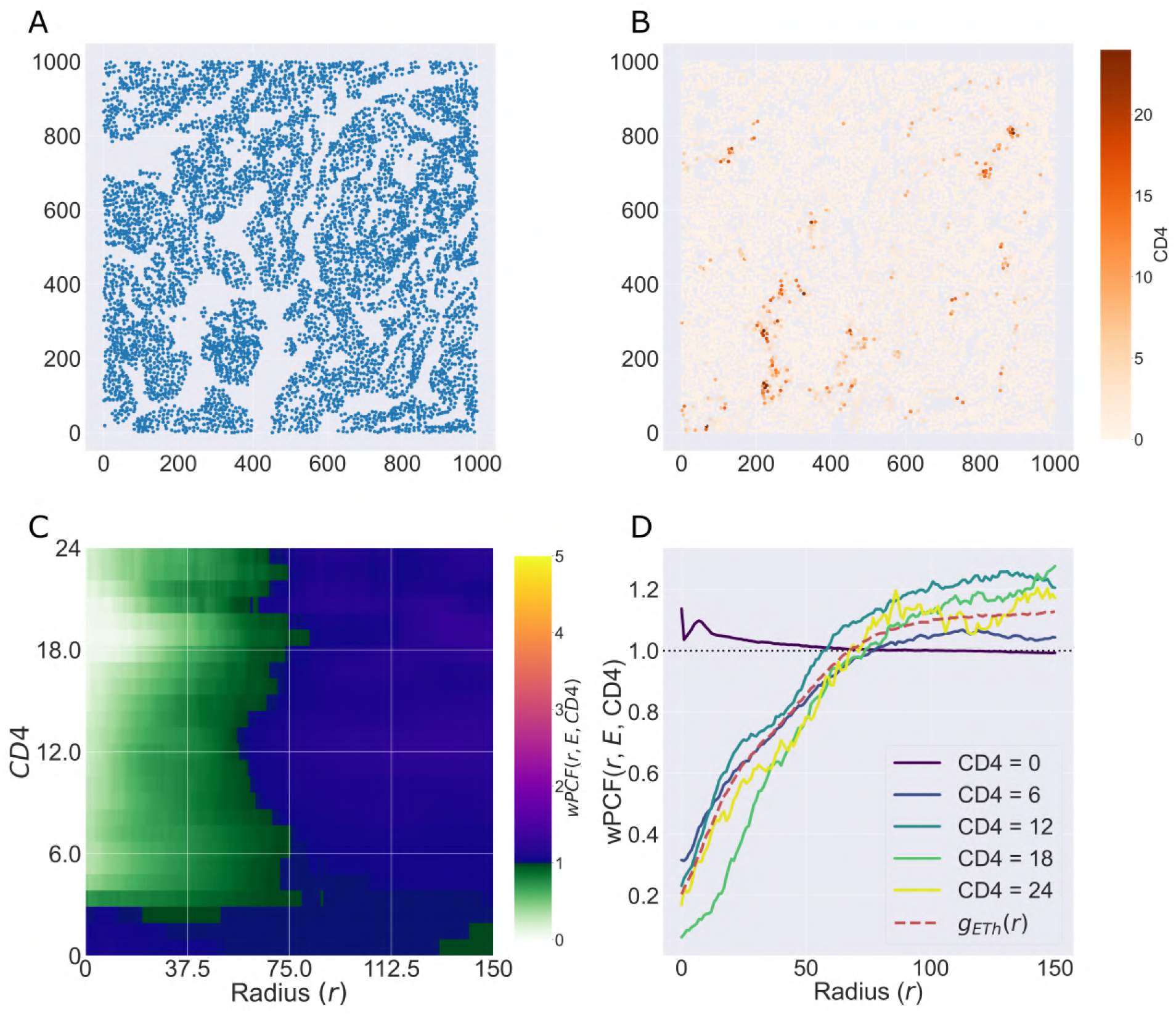
The wPCF identifies correlation between epithelial cells and cells with different CD4 expression levels. A: Epithelial cell centres. B: Cell centres labelled according to the average CD4 stain intensity within each cell. C: wPCF(*r, E*, CD4), showing clear qualitative and quantitative differences in colocalisation with epithelial cells as CD4 expression levels vary. D: Cross-sections of the wPCF in panel C. Points with low CD4 expression have a different pattern of correlation than those with higher expression. The profile for cells with high CD4 expression corresponds to the cross-PCF *g*_*ThE*_ (*r*), calculated for cells which have been manually classified as T helper cells (red dashed line). Cells with low CD4 intensity colocalise with epithelial cells, likely due to many epithelial cells having low CD4 expression. Cells with higher expression of CD4 are anti-correlated with epithelial cells for 0 ≤ *r* ⪅ 75.

Finally, in Figure 10 we show that application of the wPCF to multiplex images, without cell segmentation or classification, can identify spatial correlation. Panels A and B show points from a regular lattice sampled from the multiplex image of the ROI, at a resolution of 1 point every 5*μ*m. In panel A points are labelled according to a thresholded value of the epithelial cell marker, while in panel B they are labelled according to the CD4 intensity at that pixel. The wPCF which compares these marks is shown in panel C, and is in good qualitative and quantitative agreement with the wPCF from Figure 9 (with Δ*M* = 20 in Equation (11)). We conclude that applying the wPCF directly to pixels and stain intensities can identify the same spatial patterns of clustering and exclusion as those identified by the cross-PCF, without cell segmentation or classification.

**Figure 10:**
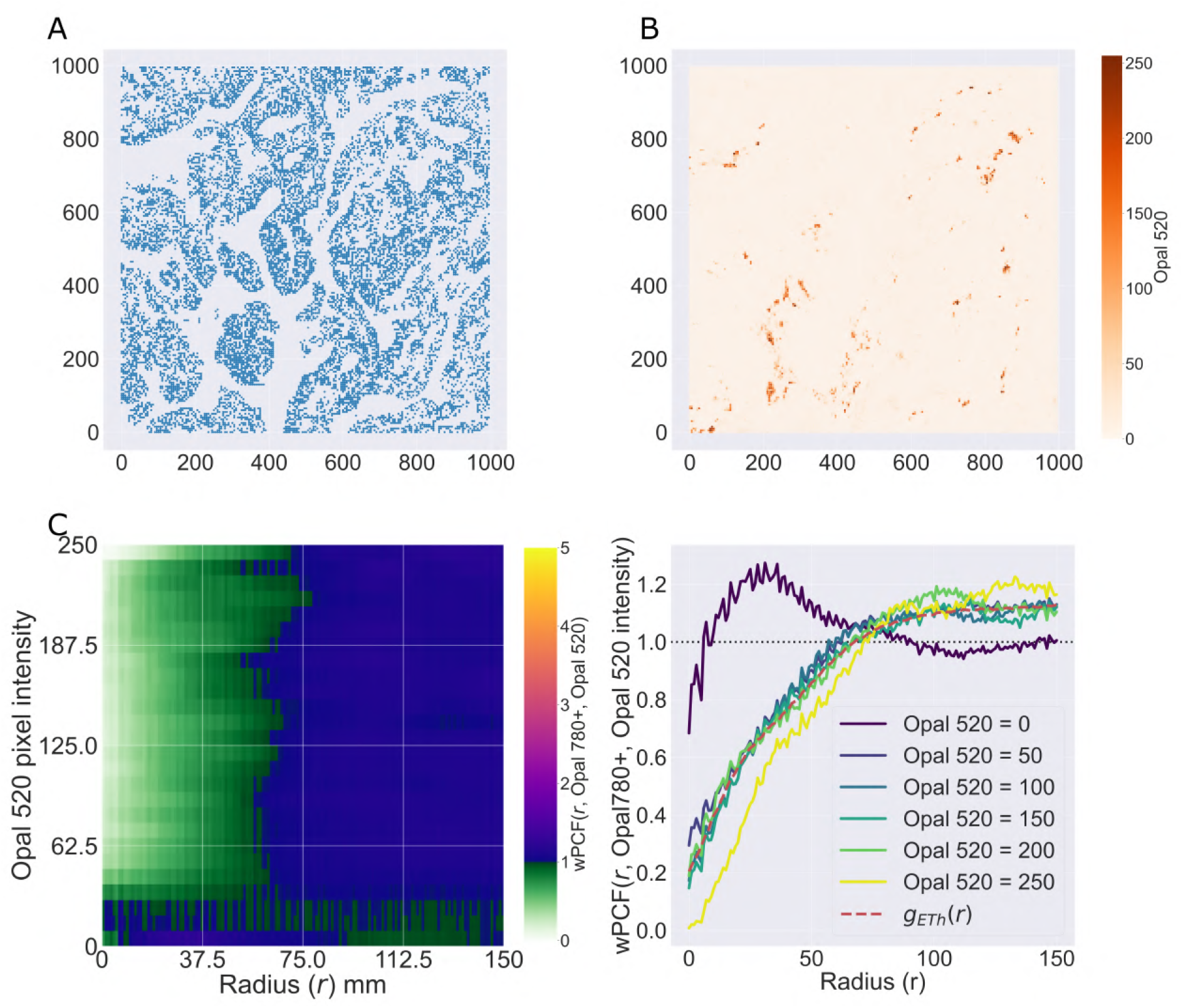
wPCF identifies correlation between epithelial cells and pixels with varying CD4 expression. The results from Figure 9 are recovered when the wPCF is calculated from points sampled from the original multiplex image using a regular 5*μ*m lattice, showing that the spatial correlation between T Helper Cells and epithelial cells can be identified without segmentation or classification. A: Pixel intensities of the Opal 520 marker (associated with CD4), sampled across the ROI on a regular 5*μ*m lattice. B: Pixels marked as Opal 780 positive (associated with epithelial cells), determined via thresholding, sampled across the ROI on a regular 5*μ*m lattice. C: wPCF describing correlation between pixels positive for Opal 780 and the pixel intensity of Opal 520. D: Cross sections of the wPCF in panel C have the same shape as the cross-PCF in panel D for pixels with high CD4 intensity

## Discussion

Multiplex images contain a wealth of spatial information, and have the potential to greatly increase the information that can be extracted from histological samples. Each image provides a high resolution map of cell locations across tissue samples which may contain millions of cells, together with detailed information about their phenotypes and morphology. As multiplex images become more widespread and as digital tools for their visualisation and analysis improve, the demand for automated methods which can extract detailed spatial information from them is increasing. Such methods should be agnostic to the technology used to generate the images, the disease under investigation, and the particular markers with which the sample has been stained.

Many existing methods can extract information from multiplex images. One popular approach involves using artificial intelligence or machine learning approaches to identify correlations between features extracted from multiplex images and clinically relevant features such as disease progression. AI methods can be extremely powerful, but are not ideally suited to all situations. In particular, an AI algorithm may require vast numbers of images for use as training data. Further, the tissue type and panel of markers chosen for staining should be consistent across the training data, thereby reducing the applicability of the algorithm to samples from different diseases (for example, an algorithm trained on multiplex images of immune cells in colorectal cancer cannot reliably be applied to images of immune cells in prostate cancer, or to images of stromal cells in colorectal cancer). AI methods can sometimes lack interpretability, making it difficult to understand which features of an image an algorithm is using and to understand when errors are likely to arise.

On the other hand, a range of statistical and mathematical methods can also describe features of multiplex images in an interpretable way. These methods may derive from a range of disciplines, such as network science, topological data analysis, and spatial statistics. They provide quantitative descriptions of specific spatial features of an image; for example, ecological analyses may describe correlations in cell counts across subregions of an ROI with a fixed area, quantifying the strength of local correlations^(45)^. Existing metrics have typically been developed to address a specific problem. As a result, multiple methods may be used to describe the same features of a point pattern. For instance, the field of spatial statistics encompasses a range of methods designed to identify correlations in point patterns, with specialised tools to address specific use cases. The PCF has been specialised to account for interactions between multiple classes of point (the cross-PCF), points generated from processes which vary across a region (inhomogeneous-PCF^(15)^), or points labelled with continuous marks (mark correlation functions, weighted-PCFs). Such metrics can provide detailed information about the spatial structure of multiplex images, even though they may have been developed for other types of data. In order to understand multiplex images using quantitative metrics, we propose the application of multiple statistics (which may derive from different mathematical fields), designed to quantify specific properties of the image.

In this paper, we have focussed on three methods for extending the PCF that have been specifically designed for application to multiplex medical images. Each is applied here for the first time to multiplex immunohistochemistry images from the Vectra Polaris system, in order to illustrate how they address limitations in the PCF. We now summarise each method in turn, focussing on their strengths and weaknesses.

### Topographical Correlation Map (TCM)

The TCM can visualise spatial correlations between pairs of cell populations across an ROI, highlighting subregions of strong positive or negative correlation that can be difficult to identify by visual inspection. Unlike the inhomogeneous-PCF, the TCM does not require prior assumptions about the homogeneity of the point process from which points are derived; rather, it identifies subregions in which local interactions between the point patterns differ from those that would be obtained under complete spatial randomness. While we have used the TCM for visualisation, it generates quantitative information which can be used for subsequent analysis. For example, the number and size of the local minima and maxima could be used as summary statistics to compare and classify images. The TCM can also be analysed via a sub/super level-set filtration^(51)^. This method from Topological Data Analysis can quantify spatial heterogeneity in heatmaps.

We note that, by design, the TCM is asymmetric (i.e., Γ_*ab*_(*r*) ≠ Γ_*ba*_(*r*)). As such, care is needed when interpreting the TCM. In particular, while Γ_*ab*_(*r*) and Γ_*ba*_(*r*) should coincide in regions of positive correlation, they may differ in regions of negative correlation. Further, regions where Γ_*ab*_(*r*) ≈ 0 can not be used to infer the presence (or absence) of either cell type without consideration of other metrics (e.g., local cell densities).

### Neighbourhood Correlation Function (NCF)

The NCF identifies whether groups of three or more cells are found in a circular neighbourhood of radius *r* more or less frequently than expected under CSR. Since it requires distance calculations between *n* cell types, the computational complexity of the NCF is at least *O*(*N*^*n*^). This limits its potential application to whole-slide images or to identifying correlations between a large number of cell types simultaneously: although calculating each enclosing circle is fast^(27)^, as the maximum number of circles which must be calculated is *N*_1_ × … × *N*_*n*_ (where *N*_*i*_ is the number of cells of type *i*) the computational effort involved increases rapidly as the total number of cells and number of cell types increase. As for the PCF, the runtime performance of the algorithm can be improved by calculating the NCF up to a maximal neighbourhood size of interest *r*; this reduces the number of *n*-wise maximum enclosing circles that must be calculated (any combination of points containing a pair separated by more than the lengthscale of interest can be immediately discarded). The NCF also relies on repeated sampling of random data to identify the expected number of neighbourhoods that would be observed under CSR. For a given region this probability can be calculated in advance to an arbitrary level of precision, and becomes more accurate with more samples. However, more research is needed to determine the minimum number of samples needed to achieve a given accuracy.

### Weighted PCF (wPCF)

The wPCF generalises the cross-PCF to data with continuous labels (e.g., cell centres which have not been classified into discrete categories, or to pixels which have not been segmented to find cell centres). As such it can be calculated without classification or cell segmentation pre-processing steps. However, this also increases the number of ‘parameters’ required. In particular, the choice of weighting function determines the ratio of signal/noise identified by the wPCF and must be considered in advance (see^(26)^ for a detailed examination of the impact of varying the weighting function on the wPCF). The tuning parameters used to construct the wPCF are in some ways similar to those used to perform cell classification (e.g. threshold values for stain intensities).

There is considerable scope for developing approaches to interpret the wPCF. The heatmap which it generates can be analysed using techniques similar to those discussed for the TCM above.

It is also possible to use the outputs from the wPCF to create a vectorised ‘spatial signature’ which can be used to cluster regions which have similar spatial structures^(26)^. Such an approach could be used to automatically identify regions with similar spatial cellular interactions, or which contain spatial patterns associated with, for example, cancer progression or disease severity. Indeed, by vectorising the spatial descriptors described within this paper the approach described in^(26)^ to identify such ‘spatial biomarkers’ could be extended.

## Conclusions

Multiplex images contain vast amounts of spatial information which can be exploited using quantitative techniques. The spatial statistics considered in this paper represent one approach to analysing this data, and benchmarking studies that compare the efficiency and insight of different methods are needed.

The methods described in this paper were designed to exploit the spatial information contained in multiplex images. We note, however, that they can be applied to multiple imaging modalities and multiple diseases. Equally, each method can be applied to generic point cloud data from contexts outside of biology.

We have previously shown that combining spatial statistics can generate more comprehensive descriptions of point data than individual metrics alone^(26,50)^. In future work, we will determine how complementary methods from mathematical fields such as spatial statistics, network science, and topology, can build upon this to provide a rigorous quantitative description of how data is spatially distributed.

## Data Availability Statement

Python code for reproducing the results described in this paper, along with the relevant data, is available on github at https://github.com/JABull1066/ExtendedCorrelationFunctions.

## Ethical Standards

The research meets all ethical guidelines, including adherence to the legal requirements of the study country.

## Author Contributions

Conceptualization: JAB. Methodology: JAB, EJM, SJL, HMB. Data curation: JAB, EJM. Data visualisation: JAB, EJM. Investigation: JAB, EJM, SJL, HMB. Writing original draft: JAB, EJM, SJL, HMB. All authors approved the final submitted draft.

## Supplementary Material

Additional supplementary material has been provided alongside this manuscript.

