## Supporting Information for "Extended correlation functions for spatial analysis of multiplex imaging data"

### S1 Analysis of other regions of interest

In this section we present supplementary analyses (without additional commentary) for three additional 1mm $\times$ 1mm regions of interest from within the same KPN mouse presented in the main text. The same cross-PCFs, NCFs, TCMs and wPCFs as those found in the main text are presented below, and show the same general results presented in the main text.

| Celltype | Marker | Region 1 - $n$ | Region 2 - $n$ | Region 3 - $n$ |
|---|---|---|---|---|
| Epithelium | E-Cadherin | 5122 | 5812 | 5737 |
| Macrophage | CD68 | 549 | 287 | 467 |
| T helper cell | CD4+ FoxP3- | 965 | 91 | 666 |
| Neutrophil | Ly6G | 116 | 165 | 163 |
| Cytotoxic T cell | CD8 | 163 | 14 | 130 |
| Regulatory T cell | CD4+ FoxP3+ | 25 | 5 | 49 |

Table 1: Number of cells, $n$, for each supplementary region

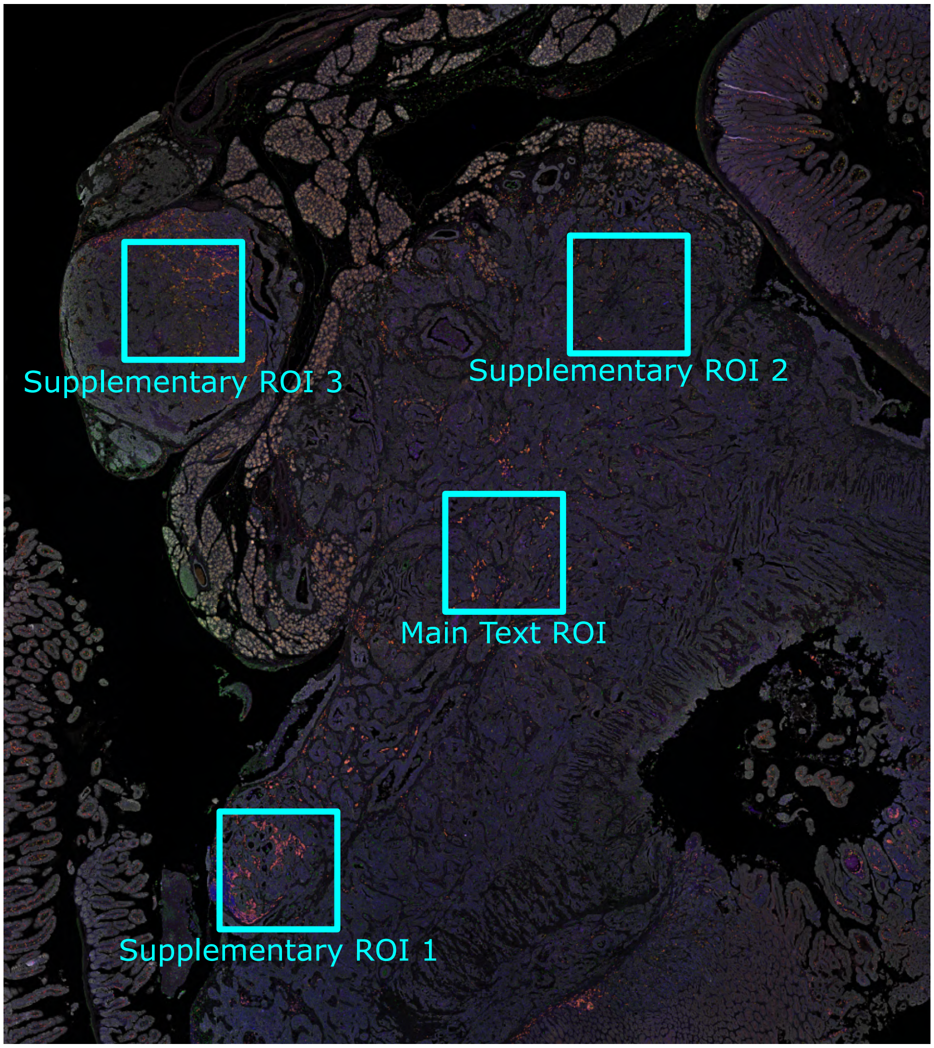


Figure S1: Location of 1mm × 1mm ROIs within the wider tissue context

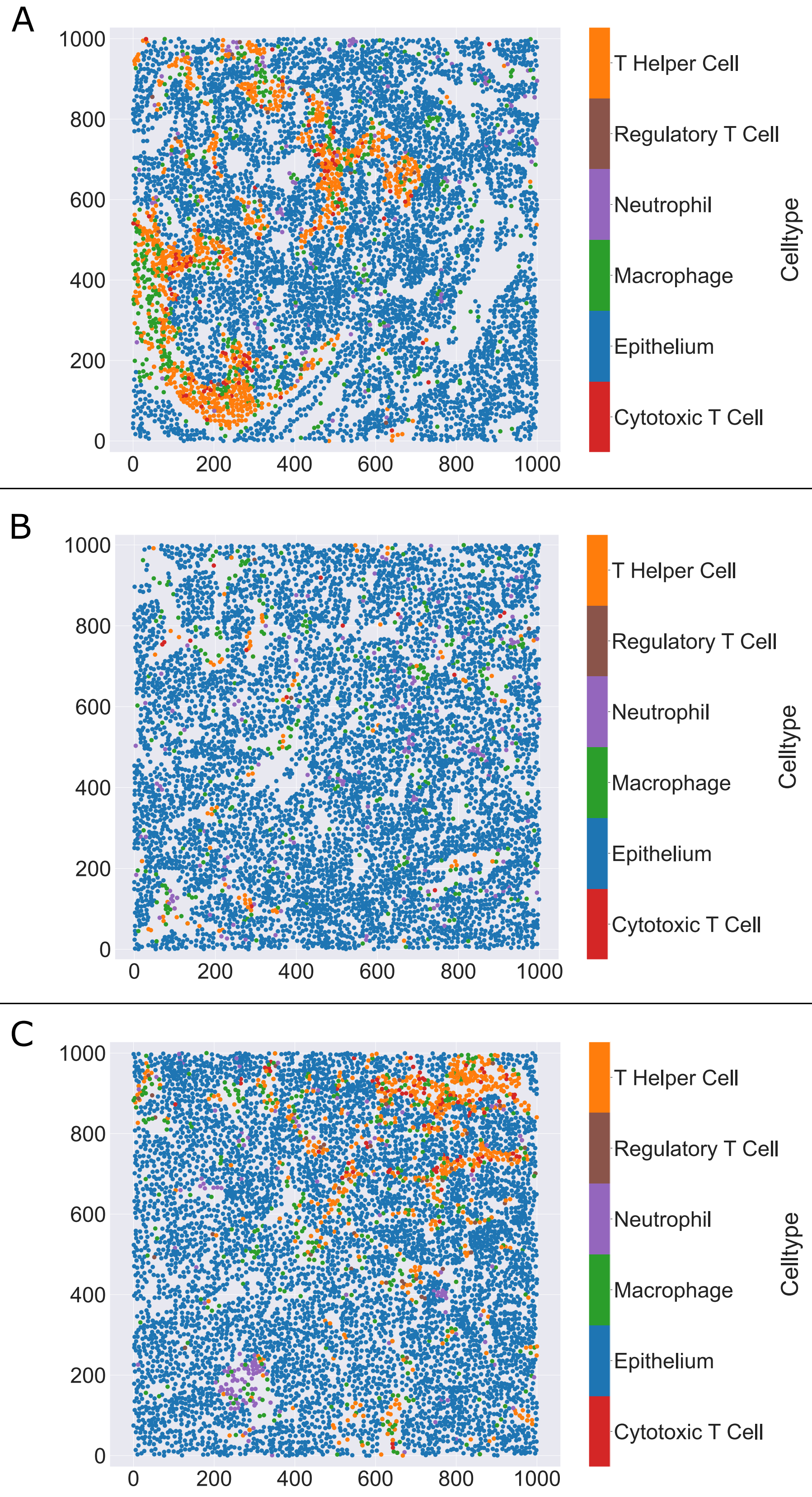


Figure S2: Cell centre locations for supplementary ROIs 1 (A), 2 (B) and 3 (C)

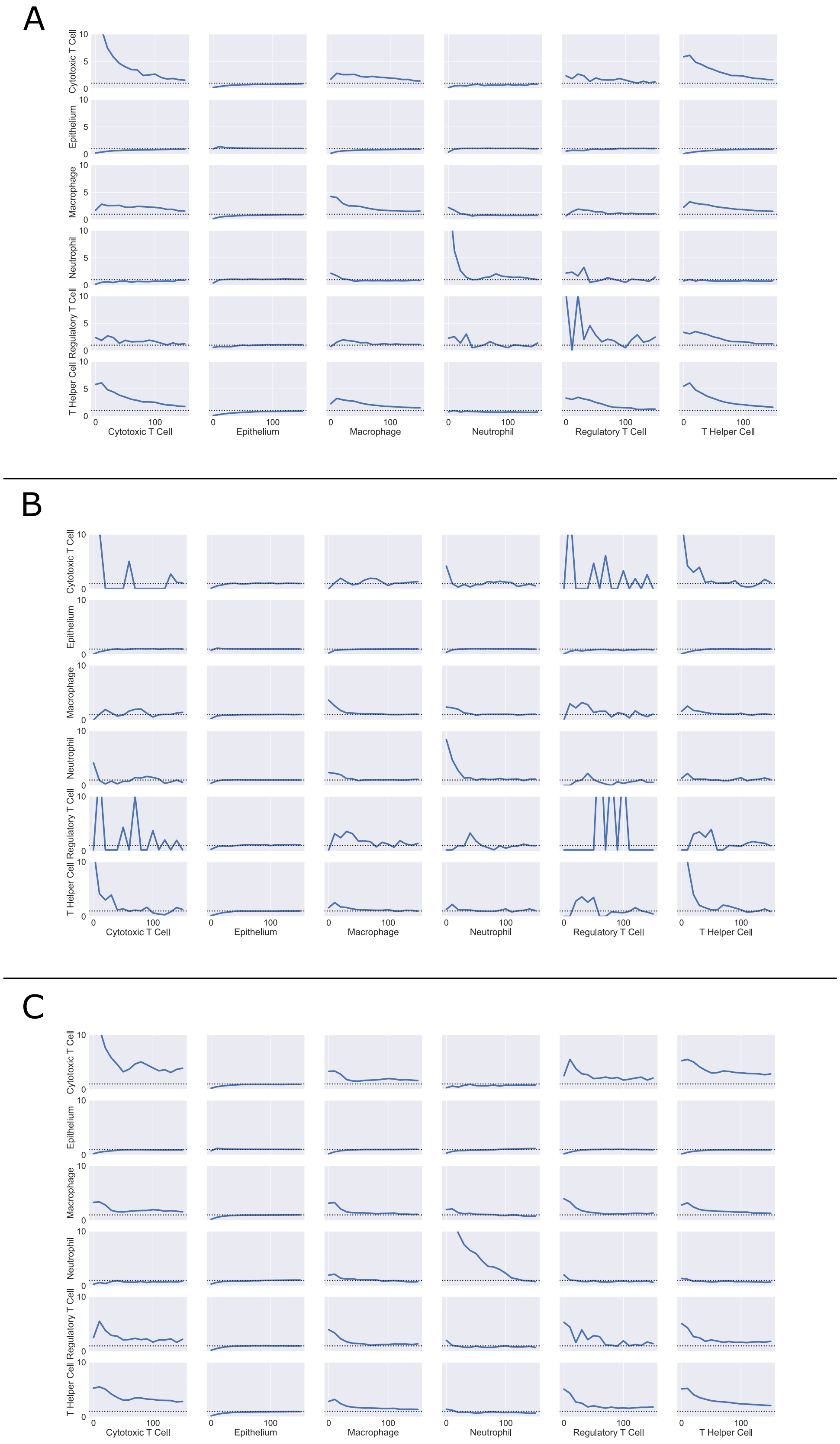


Figure S3: All cross-PCFs from supplementary ROIs 1 (A), 2 (B) and 3 (C)

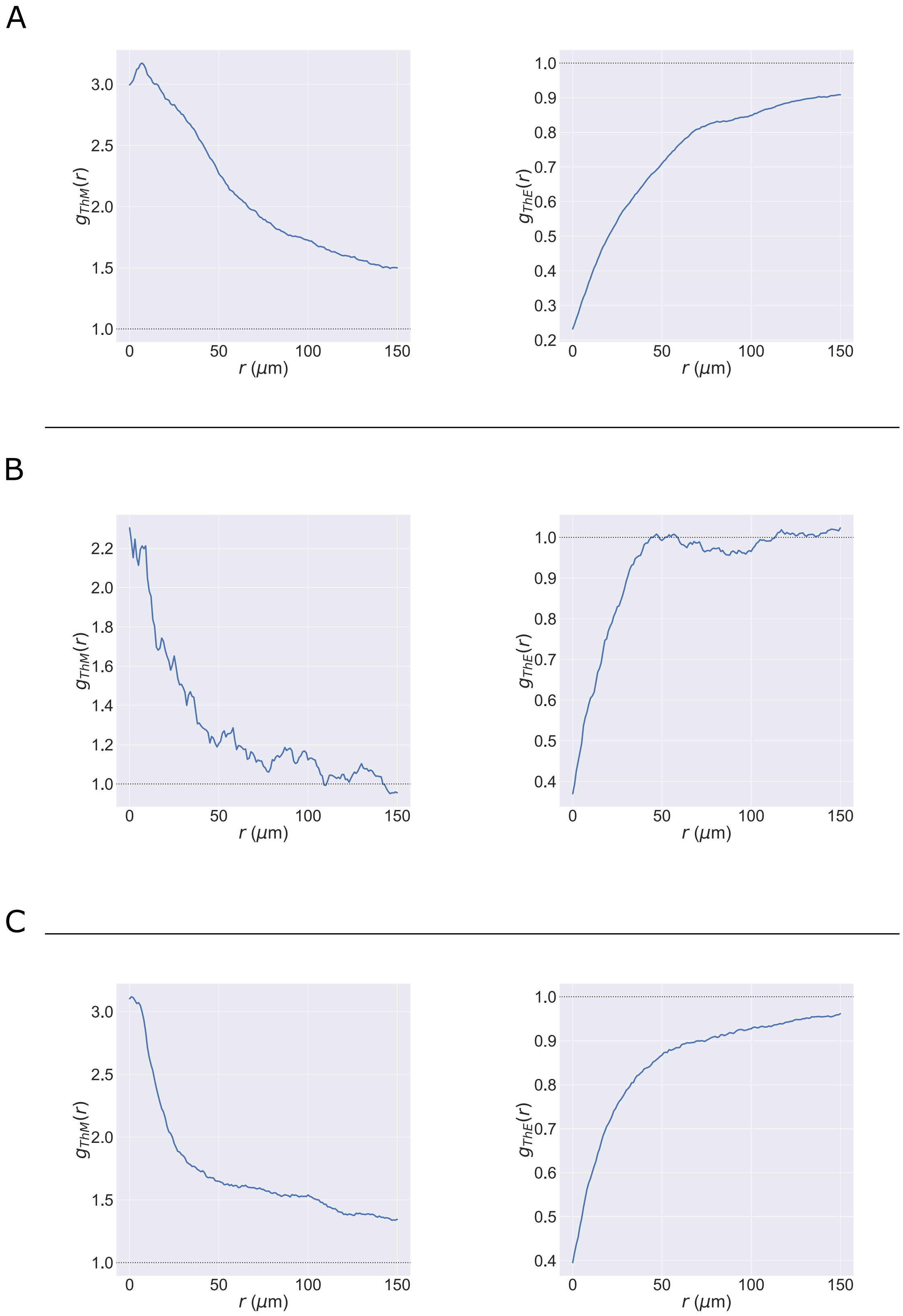




Figure S4: Cross-PCFs between T Helper Cells and Macrophages (left) / Epithelium (right) from supplementary ROIs 1 (A), 2 (B) and 3 (C)

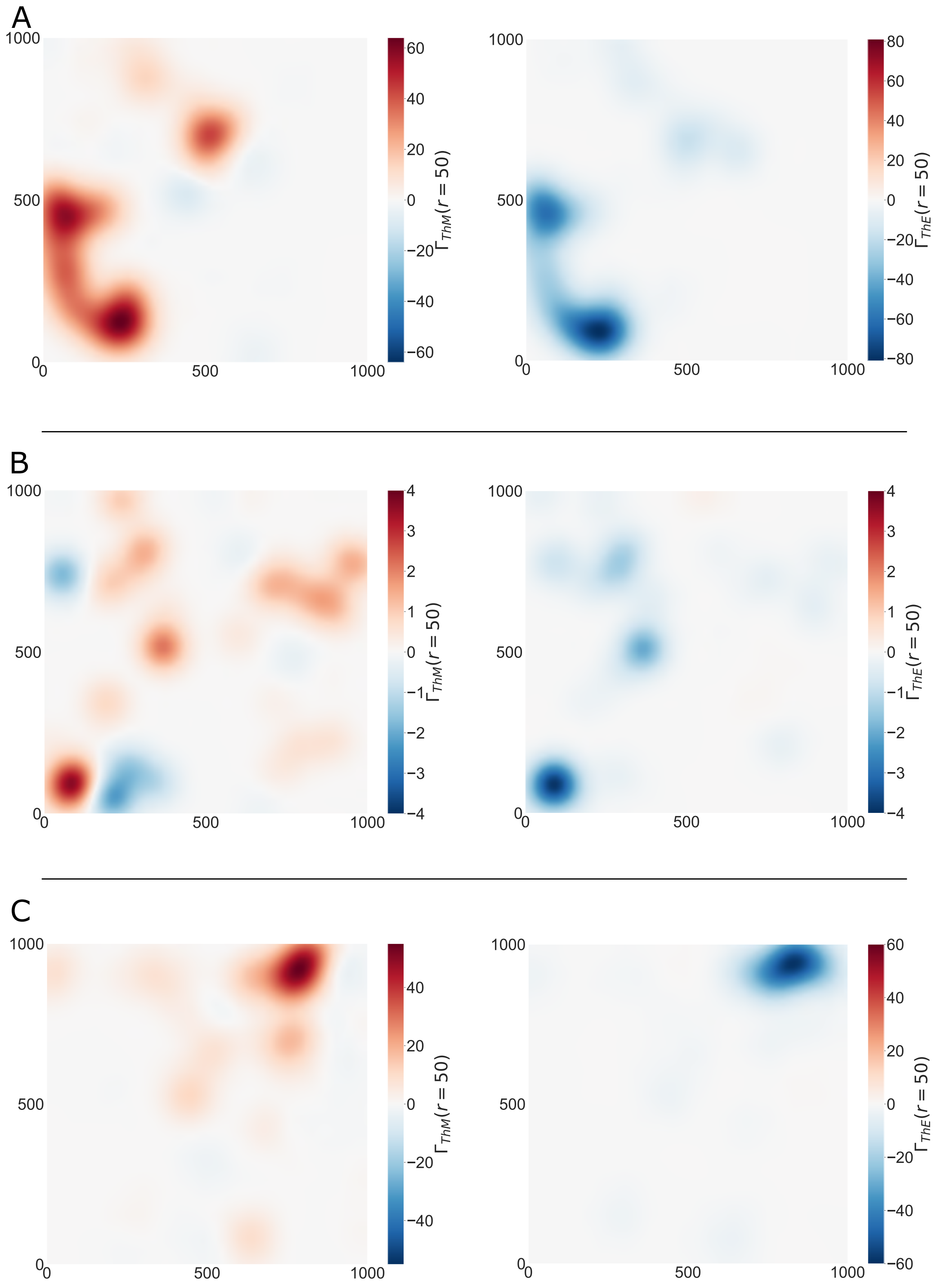




Figure S5: TCMs between T Helper Cells and Macrophages (left) / Epithelium (right) from supplementary ROIs 1 (A), 2 (B) and 3 (C)

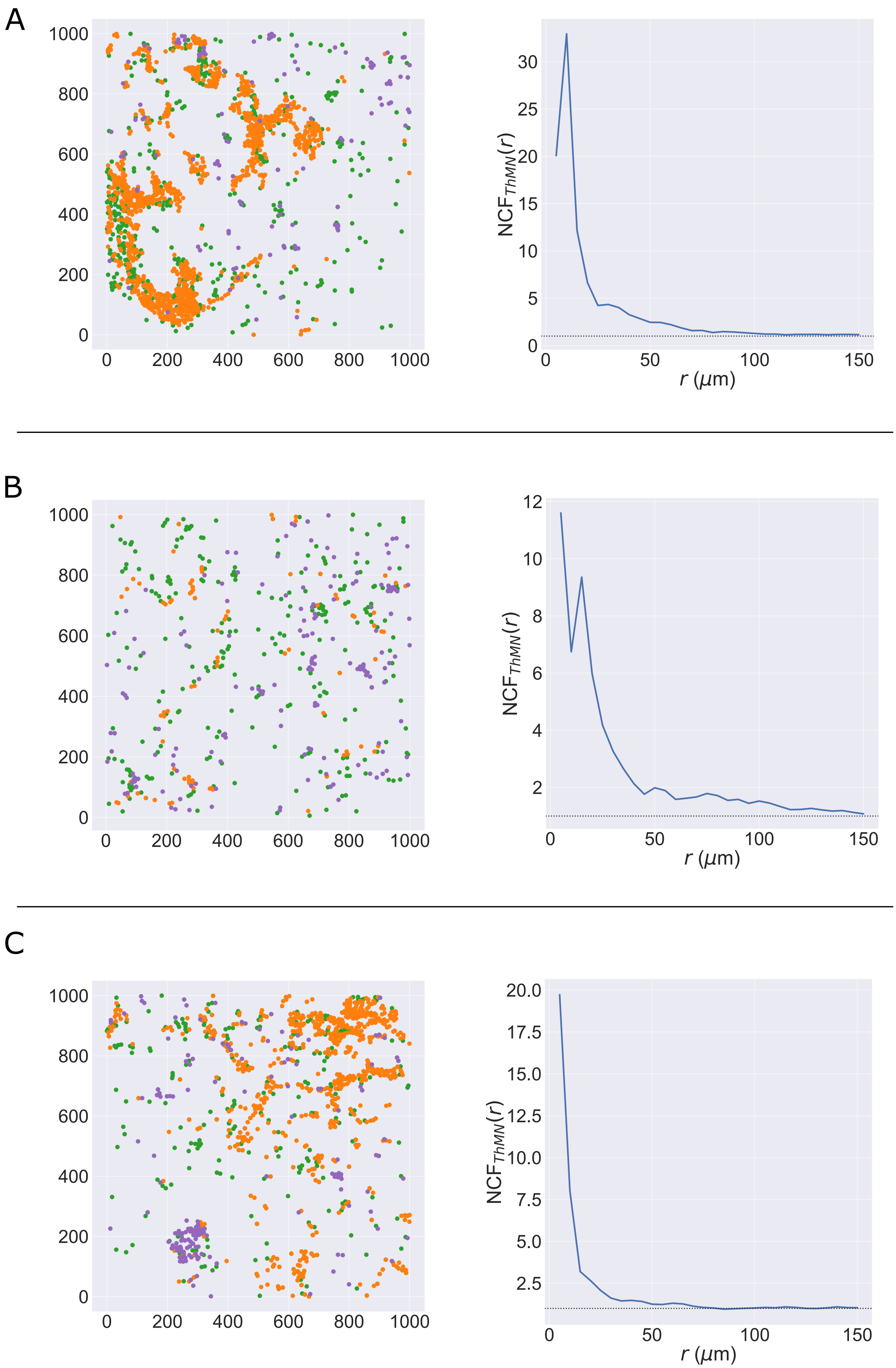




Figure S6: Left: locations of T Helper cells (orange), Macrophages (green) and Neutrophils (purple) from supplementary ROIs 1 (A), 2 (B) and 3 (C). Right: NCF between T Helper cells, Macrophages and Neutrophils from supplementary ROIs 1 (A), 2 (B) and 3 (C).

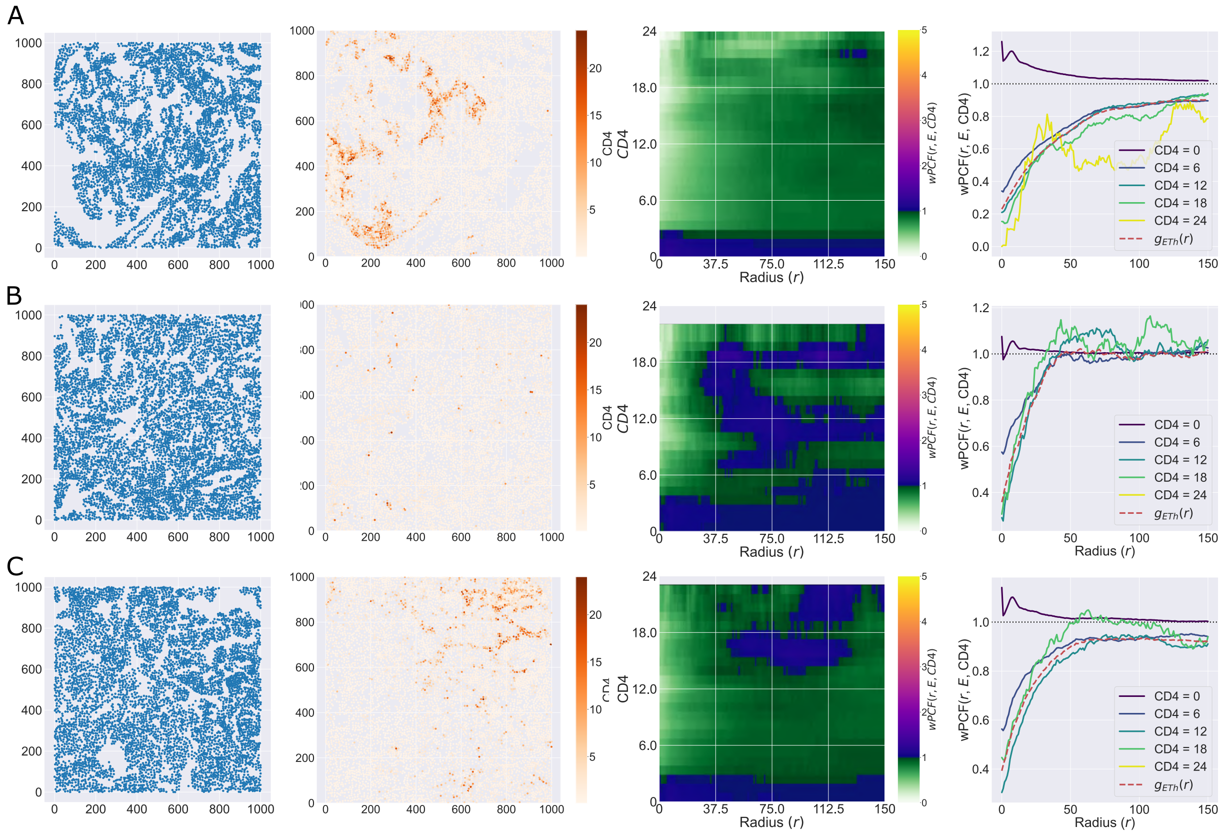

Figure S7: From left: i) locations of epithelial (cancer) cells; ii) all cells labelled according to CD4 intensity; iii) wPCF($r$, $E$, CD4); iv) cross-sections of wPCF($r$, $E$, CD4) at fixed values for the CD4 intensity. From supplementary ROIs 1 (A), 2 (B) and 3 (C).
